# Exostosin-1 Glycosyltransferase Regulates Endoplasmic Reticulum Architecture and Dynamics

**DOI:** 10.1101/2020.09.02.275925

**Authors:** Despoina Kerselidou, Bushra Saeed Dohai, David R. Nelson, Sarah Daakour, Nicolas De Cock, Dae-Kyum Kim, Julien Olivet, Diana C. El Assal, Ashish Jaiswal, Deeya Saha, Charlotte Pain, Filip Matthijssens, Pierre Lemaitre, Michael Herfs, Julien Chapuis, Bart Ghesquiere, Didier Vertommen, Verena Kriechbaumer, Kèvin Knoops, Carmen Lopez-Iglesias, Marc van Zandvoort, Jean-Charles Lambert, Julien Hanson, Christophe Desmet, Marc Thiry, Kyle J. Lauersen, Marc Vidal, Pieter Van Vlierberghe, Franck Dequiedt, Kourosh Salehi-Ashtiani, Jean-Claude Twizere

## Abstract

The endoplasmic reticulum (ER) is a central eukaryotic organelle with a tubular network made of hairpin proteins linked by hydrolysis of GTP nucleotides. Among post-translational modifications initiated at the ER level, glycosylation is the most common reaction. However, our understanding of the impact of glycosylation on ER structure remains unclear. Here, we show that Exostosin-1 (EXT1) glycosyltransferase, an enzyme involved in *N*-glycosylation, is a key regulator of ER morphology and dynamics. We have integrated multi-omics data and super-resolution imaging to characterize the broad effect of EXT1 inactivation, including ER shape-dynamics-function relationships in mammalian cells. We have observed that, inactivating EXT1 induces cell enlargement and enhances metabolic switches such as protein secretion. In particular, suppressing EXT1 in mouse thymocytes causes developmental dysfunctions associated to ER network extension. Our findings suggest that EXT1 drives glycosylation reactions involving ER structural proteins and high-energy nucleotide sugars, which might also apply to other organelles.

## INTRODUCTION

The ER is one of the largest organelles of eukaryotic cells (Porter et al., 1945). It facilitates communication with other intracellular organelles through its connection to the nuclear envelope and regulates interactions with the external environment *via* the secretion of proteins, polysaccharides, and lipids (Phillips and Voeltz, 2016).

The ER is involved in numerous cellular processes, from lipid turnover to protein secretion and glycosylation. Part of the metabolic flexibility of the ER is mediated by its dynamic and adaptable shape (Palade and Porter, 1954; Porter and Palade, 1957). During normal cell homeostasis, the ER is a complex network of tubules and flat matrices that are in continuous motion. These sub-structures form a three-dimensional, regularly shaped network, which derives its form from the lipid bilayer and different groups of membrane-associated proteins (Terasaki et al., 2013). High-curvature regions, such as ER tubules and edges of ER sheets, are built by the oligomerization of hydrophobic hairpin domain-containing reticulons (RTNs), and receptor expression enhancing proteins (REEPs) (Voeltz et al., 2006; Yang and Strittmatter, 2007). Flat matrices are formed by atlastin (ATL) GTPase-ER membrane associations; these proteins dimerize in opposing layers to hold lipid bilayers in this conformation (Liu et al., 2015). The curvature of flat matrices is mediated by the luminal bridging cytoskeleton-linking membrane protein 63 (CLIMP63) (Shibata et al., 2010).

It is currently believed that cooperation between ER shape and luminal dynamics dictates ER functions (Schwarz and Blower, 2016). ER sheets are the primary sites for translation, translocation, and folding of integral membrane-bound as well as secreted proteins, while ER tubules are thought to be involved in other ER functions such as lipid synthesis and interactions with other organelles (Shibata et al., 2006; Voeltz et al., 2002). Cells actively adapt their ER tubules/sheets balance and dynamics to coordinate its morphology and function, in accordance with cellular demands (Westrate et al., 2015). However, the molecular mechanisms underlying overall maintenance and flexibility of the ER network remain poorly characterized.

One of the key roles the ER plays for the cell is the secretion of extracellular products and glycosylation of proteins. Our understanding of the mechanisms of glycosylation are now advanced enough to engineer cell lines to confer customized glyco motifs for biotechnological and medicinal applications. Yet, our understanding of how these motifs affect the intracellular dynamics of cellular homeostasis is limited. Glycosylation proceeds by the synthesis of glycans and attachment to the acceptor peptide, which is initiated in the ER and terminates in the Golgi apparatus (Reily et al., 2019). Glycosylation is well known to regulate the physical properties of different glycolipid and glycoprotein biopolymers at the surface of mammalian cells by controlling plasma membrane and cell coat morphologies (Shurer et al., 2019). The impact of intracellular ER membrane protein component glycosylation, their interactions with membranes, and their contributions to cellular dynamics are entirely unknown. While permanent interactions between membrane curvature proteins are sufficient to form the basic ER structure (Powers et al., 2017; Schwarz and Blower, 2016), protein-protein interactions and post-translational modifications (PTMs) may participate in its dynamic shaping. It is, therefore, essential to decipher how glycosylation affects ER dynamics within the cell in addition to defining the sequential steps leading to final glycosylation species of extracellular proteins.

Synthesis of glycans and their attachment to proteins occurs by the sequential activities of glycosyltransferases and glycosidases that compete for activated glycans and overlapping substrates (Reily et al., 2019). Protein *N*-glycosylation occurs in the ER lumen and is catalyzed by the oligosaccharyltransferase complex (OST) which is composed of eight proteins in metazoans (ribophorin, DAD1, Tusc3, OST4, TMEM258, OST48, and catalytic subunits STT3A and STT3B) (Kornfeld and Kornfeld, 1985). The final composition of oligosaccharide chains bound to glycoproteins depends not only on localization and abundance of these enzymes but also on the availability and heterogeneity of sugar substrates. Exostosin-1 (EXT1) is an ER resident glycosyltransferase involved in the polymerization of heparan sulfate (HS) (McCormick et al., 1998, 1999). HS molecules are found in all animal tissues and play a key role in many biological activities, including development and cancer. While investigating the role of EXT1 in thymocytes development and cancer, we discovered that reduction of the EXT1 protein results in global changes in cellular homeostasis, including cell size, organelle shapes and interactions, and cellular metabolism. We show that reprogramming of glycan moieties by reduction of this regulator can profoundly change ER structure concurrent with global metabolic shifts in protein and membrane lipid synthesis in cells.

## RESULTS

### Developmental defects following EXT1 inactivation are mediated by its genetic interactions

In a systematic interactome study, we previously showed that EXT1, an ER-resident type II transmembrane glycosyltransferase, interacts with Notch1, a type I transmembrane receptor that is frequently mutated in cancers (Daakour et al., 2016). Notch1 is essential for the development of numerous cell types, including thymocytes (Artavanis-Tsakonas et al., 1995; Wolfer et al., 2002). We thus hypothesized that EXT1 might play a physiological role in T cell development, potentially associated with its glycosyltransferase function and ER residency (Figure 1A). Because homozygous *EXT1* null mice die at embryonic day 8.5 (Lin et al., 2009), we crossed *EXT1*^*F/F*^ (Inatani et al., 2003) mice with mice expressing the Cre recombinase under the control of *lck* proximal promoter (Lee et al., 2001) to specifically target *EXT1* in early developing thymocytes. We found that *EXT1* inactivation affects the early stage of thymocyte development, with a significant accumulation of immature double negative CD4^-^, CD8^-^ cells (DN) (p<0.01, Figure 1B-C). We also used a *Notch*^*F/F*^ line (Radtke et al., 1999) to generate conditional knockout of *Notch1*, or both *EXT1 and Notch1* genes simultaneously (Figure S1A-B). As previously shown (Wolfer et al., 2002), *Notch1* inactivation leads to the accumulation of DN, at a higher extend compared to *EXT1 k*.*o*. (p<0.0001, Figure 1B-C). In both cases, *Notch1* or *EXT1* inactivation affect the late stages of thymocytes developmental stages DN3 and DN4 (Figure 1D-E). Surprisingly, transgenic mice with thymic inactivation of both genes exhibit a normal phenotype, suggesting a genetic suppression interaction (GSI) between *Notch1* and *EXT1* in thymocytes (Figure 1B-E). Because the altered phenotype in *Notch1* deficient thymocytes was rescued by *EXT1* knockout, we concluded that *EXT1* may act as a functional suppressor partner of the *Notch1* receptor *in vivo*.

**Figure 1.**
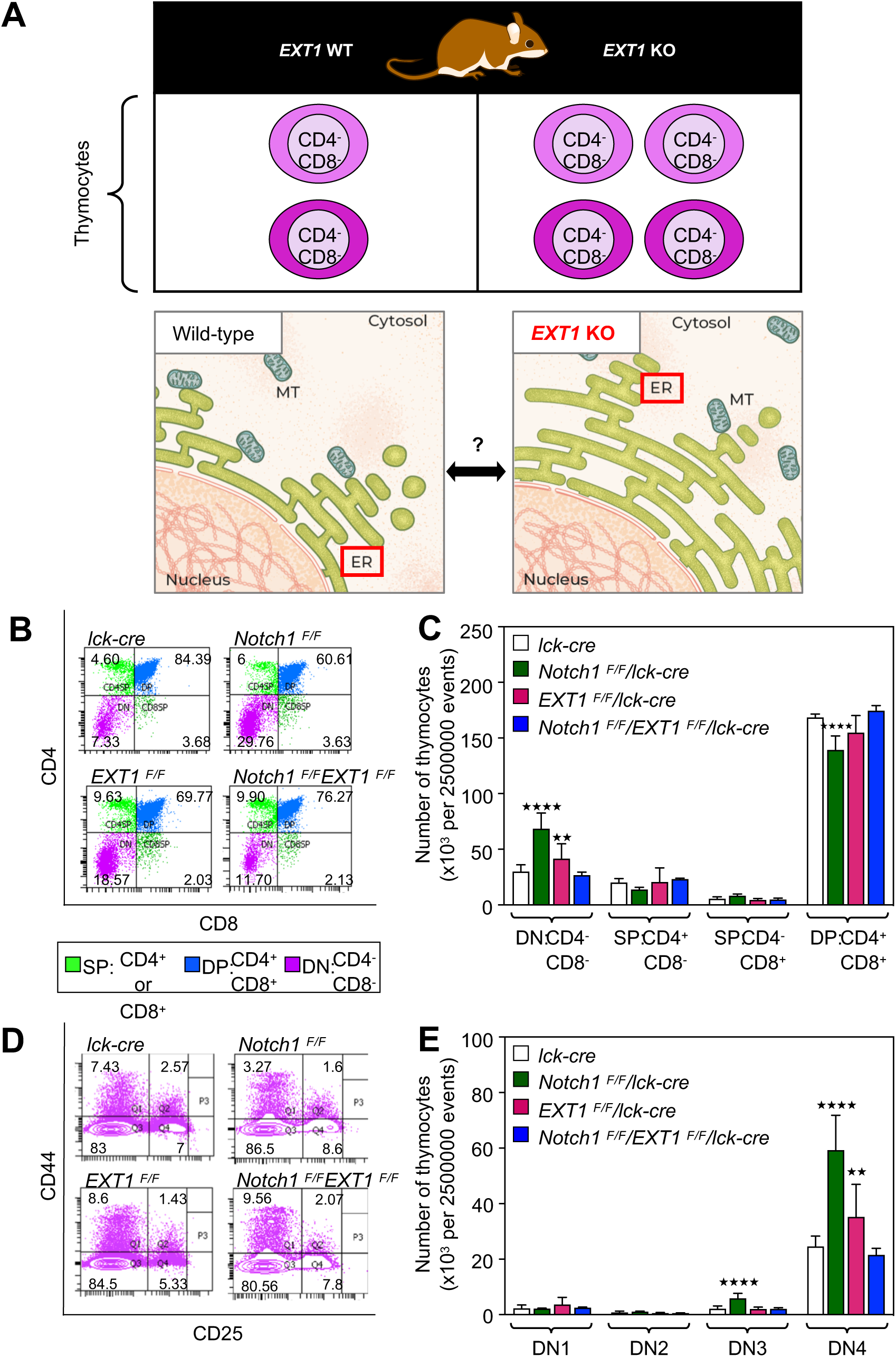
Developmental defects following EXT1 inactivation are mediated by its genetic interactions. **(A)** Framework to study the role of ER-resident EXT1 in thymocytes development. **(B)** Representative FACS plots showing surface phenotype of CD4 and CD8 T-cells in thymocytes. Cell percentages are shown in quadrants. **(C)** The absolute number of thymocytes (out of 2500000 total events) showing surface receptor expression of DN, SP and DP populations. n = 6 mice (*lck-cre, Notch1*^*F/F*^*/lck-cre, EXT1*^*F/F*^*/lck-cre, Notch1*^*F/F*^*/EXT1*^*F/F*^*/lck-cre*). **(D)** Representative FACS plots showing surface expression of CD44 and CD25 markers in DN populations. Cell percentages are shown in quadrants. **(E)** The absolute number of DN1, DN2, DN3 and DN4 cells (out of 2500000 total events).

### Cancer dependency to EXT1 expression is associated with perturbations of ER structures

The unexpected phenotype generated by *Notch1* and *EXT1* double *k*.*o*. allowed us to hypothesize that, *EXT1* could be a candidate synthetic lethal (SL) (Feng et al., 2019; Lee et al., 2018b) or synthetic dosage lethal (SDL) (Megchelenbrink et al., 2015) gene with activated oncogenic Notch1. To test the SDL hypothesis, we knocked down or overexpressed EXT1 in Jurkat, a T-cell acute lymphoblastic leukemia (ALL) cell line, which have altered Notch1 signaling (Figure S1C-D). Dosage variations did not influence Jurkat cell proliferation (Figure S1E). However, when injected into non-obese diabetic/severe combined immunodeficiency (NOD/SCID) mice (van der Loo et al., 1998), we observed a striking and significant reduction in tumorigenicity following EXT1 knockdown (p<0.0001, Figure 2A-B). Concurrently, overexpression of EXT1 was found to cause more tumor burden than control Jurkat T-ALL cells (Figure 2C-D), demonstrating a dosage lethality effect of EXT1 in the Jurkat T-cell model. To test the SL hypothesis, we interrogated the gold standard SL gene pairs across different cancer types (Lee et al., 2018b). In these cancer patient cell lines, we did not observe any SL interaction between *EXT1* and *Notch1*, or *EXT1* and the Notch1 ubiquitin ligase encoding gene *FBXW7*. However, *EXT1* and *Notch1* do synthetically interact with several shared genes, including important oncogenes: *KRAS, PTEN, BRCAC2*, and *MYC* (Figure S1F). *EXT1* also appears as a clinically significant hub, for which downregulation by shRNA presented numerous SL interactions relevant for various cancer types (Figure S1F). An exploration of the cancer genome atlas, TCGA, (Ghandi et al., 2019; Iorio et al., 2016; McDonald et al., 2017) for somatic mutations in different cancer cell lines and tumors also highlighted *EXT1* as a clinically relevant hub (Figures S1G-H). These findings suggest that *EXT1* is a genetic suppressor of *Notch1* and a potential precision therapeutic target in cancers for which Notch1 and other selected oncogenes are activated.

**Figure 2.**
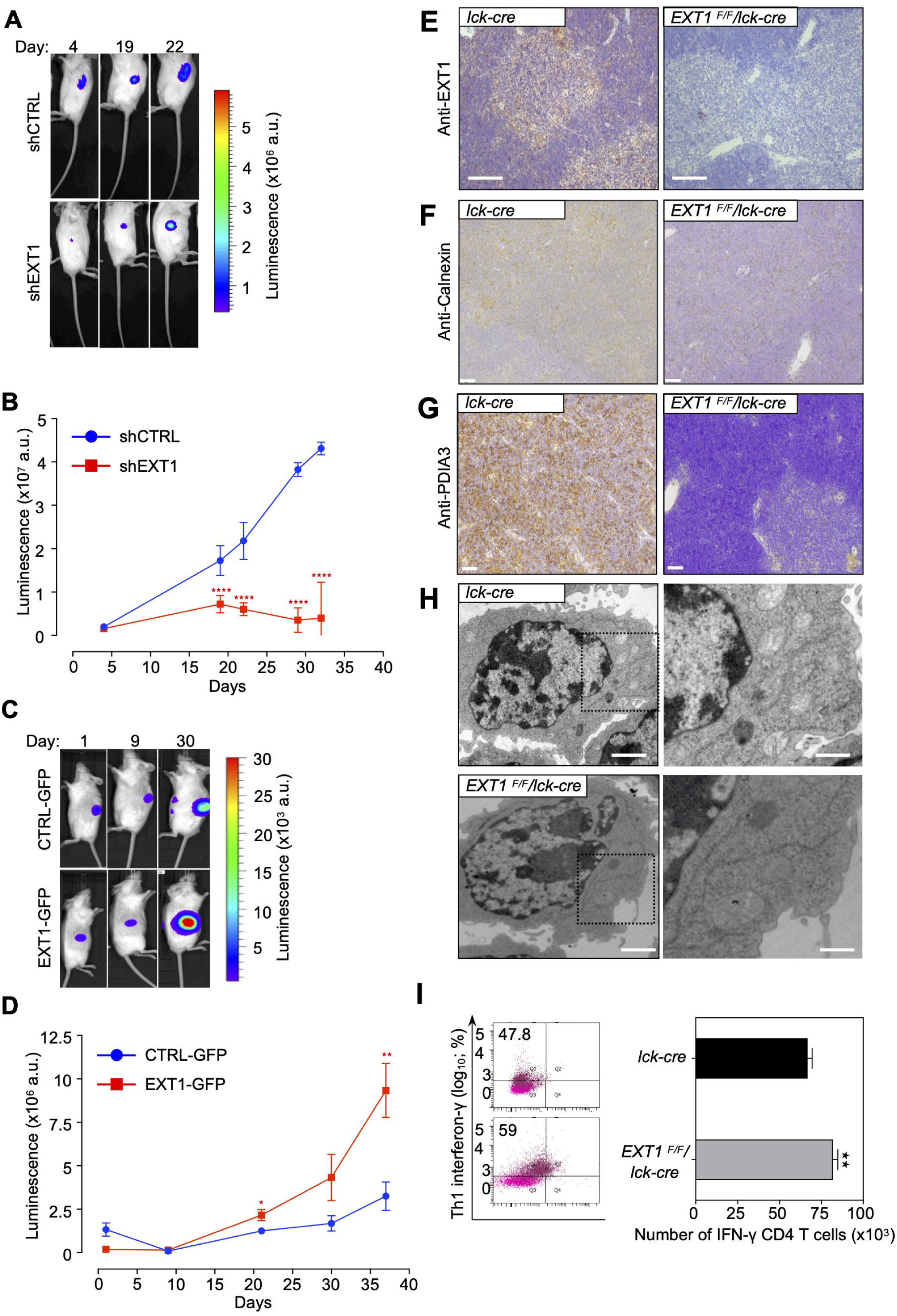
Cancer dependency to EXT1 expression is associated with perturbations of ER structures. **(A-B)** Follow-up of the tumor progression *via* bioluminescence after injection of 2⨯10^6^ control and shEXT1 Jurkat cells at opposites sites of NOD/SCID mice. **(C-D)** As in A-B for CTRL-GFP and EXT1-GFP cells. One-way ANOVA: **p<0.01, ****p<0.0001. **(E)** Immunohistochemistry staining of EXT1 Protein in thymus of *lck-cre* (left panel) and *EXT1*^F/F^/*lck-cre* (right panel) mice. Scale bar, 2 μm **(F)** As in (E) but immunohistochemistry staining with Anti-Calnexin antibody. Scale bar, 2 μm **(G)** As in (E) but immunohistochemistry staining with Anti-PDIA3 antibody. Scale bar, 2 μm **(H)** TEM of ER in activated T-cells from murine peripheral lymph nodes and spleen. Scale bar, 2 μm. **(I)** The percentages of IFN-γ in Th1 from *lck-cre, EXT1*^F/F^/*lck-cre* mice are shown in quadrants. Percentages of IFN-γ positive cells are shown in quadrants. Bar graphs represent mean number + SD. One-way ANOVA: **p<0.01. See also Figures S1 and S2.

We next sought to investigate the molecular mechanisms underlying the identification of *EXT1* as a suppressor hub. Because the EXT1 protein localizes predominantly in the endoplasmic reticulum (ER) (McCormick et al., 1998, 1999), we first assessed ER phenotypes of thymus from mutant mice with inactivation of *EXT1* (*EXT1*^*F/F*^*/lck-cre*) (Figure 2E). Immunohistochemistry was used to examine the expression of the ER resident molecular chaperones calnexin and PDIA3, and we observed a striking reduction in the expression of both markers in structural cells of thymus from conditional *EXT1 k*.*o*. compared to control mice (Figures 2F-G). To gain information on the consequences of *EXT1 k*.*o*. in the thymus, we analyzed mature lymphocytes migrating from the thymus. Transmission electron microscopy (TEM) analysis highlighted an unusual elongated ER morphology in activated CD4+T cells from *EXT1 k*.*o*. mice, (Figure 2H), with a concomitant increase in the T-helper 1 interferon-gamma (IFN1-γ)-producing cell population (Figure 2I). The results suggest an important role of EXT1 in ER organization in T-cells.

### EXT1 downregulation causes ER extension and cell size increase

The elongated morphology of ER in *EXT1 k*.*o*. mice thymocytes was unexpected. To rule out cell line-specific effects, we assessed the ultrastructural ER morphology in HeLa, HEK293, and Jurkat cells. Following *EXT1* knockdown (k.d.), we observed a dramatic elongation of ER tubules in all cell lines, with for HeLa cell line, an average length of 109.6±25.3 µm compared to 19.0±8.0 µm in control cells (Figures 3A-B, S2A-D and Table S1). The depletion of other members of the exostosin family (*EXT2, EXTL1-3*) did not lead to similar ER changes (Figures S2E-F). Probably as a consequence of ER extension, the cell area increased by ∼2 fold in *EXT1 k*.*d*. cells compared to controls (133.9±36.8 and 68.5±12.5 μm^2^, respectively) (Figure 3C-D and Table S1). Cell size is of fundamental importance to all biological processes, and it is strictly regulated to keep a balance between growth and division (Campos et al., 2014; Turner et al., 2012). We did not observe any significant effect on proliferation following *EXT1 k*.*d*. (Figures S3A-B), suggesting that *EXT1 k*.*d*. cells might have undergone an important adaptive change of the size-threshold following ER extension and internal cellular architecture rearrangement (Figure 3A and 3C).

**Figure 3.**
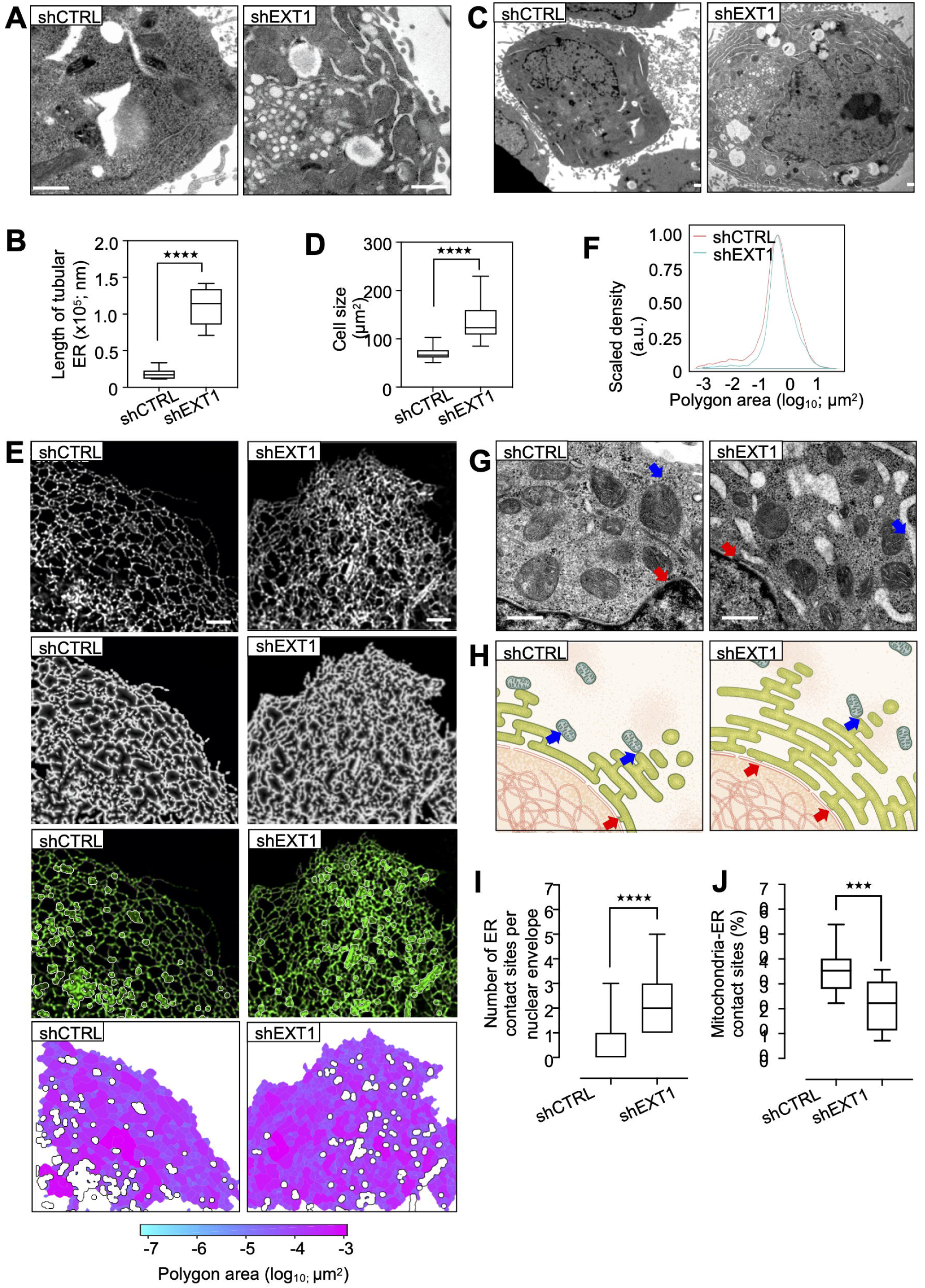
EXT1 downregulation causes ER extension and cell size increase. **(A)** TEM of the ultrastructure ER of HeLa cells. Scale bar, 2 μm. **(B)** Quantification of the length of tubular RER (nm)/cell (n = 10). Boxplot indicates the mean and whiskers show the minimum and maximum values. One-way ANOVA: ****p<0.0001. **(C)** TEM of HeLa shCTRL and shEXT1 HeLa cells Scale bar, 2 μm. **(D)** Quantification of the cell area (Table S1). **(E)** Confocal fluorescence of Cos7 cells shCTRL or shEXT1 transiently expressing mEmerald-Sec61b. From top to bottom: original image, skeleton, overlay of the skeleton (purple), the cisternae (white) and the original image (green), polygonal regions map, color-coded by size. **(F)** Quantitative analysis based on the skeletonization model of Cos7 cells expressing mEmerald-Sec61b in shCTRL and shEXT1 condition. Polygon area (log_10_) in X-axis is plotted against scaled density in Y-axis (n = 19-24). **(G)** TEM of ER-mitochondria and ER-nuclear envelope contact sites in HeLa shCTRL and shEXT1 cells. Arrows in blue and red highlight the contact sites. The nuclear envelope-ER contact sites increase and the number of mitochondria-ER contact sites decreases following EXT1 depletion. Scale bar, 500 nm. **(H)** Schematic representation of the ER-other organelle contact sites as used for the statistical analysis of the different parameters. **(I)** Quantification of the ER-nuclear envelope contact sites in boxplot indicating the mean and whiskers show the minimum and maximum values (n = 10-18). **(J)** As in (I) but percentage of ER-mitochondria contact sites (n = 10). See also Figure S3 and Table S1.

To analyze the ER luminal structural rearrangements, we quantified ER membrane structures marked with SEC61b by confocal microscopy and a segmentation algorithm that excludes insufficient fluorescent intensity. This strategy generates a single-pixel-wide network to allow quantification of individual tubule morphological features. The tubular ER network was altered in *EXT1 k*.*d*. cells, and exhibited a denser and more reticulated phenotype in comparison to controls (Figure 3E). Measurements of the polygonal area of ER tubular network in these cells were 0.778 µm^2^, which is a reduction from 0.946 µm^2^ in controls (Figure 3F). Other tubular and cisternal ER metrics were unaffected (Figures S3C-E), suggesting that the dense tubular network might relate to a more crowded ER lumen.

We next analyzed ER interactions with other organelles and counted significantly more peripheral ER-nuclear envelope (2.3±1.2 versus 0.6±0.9) and less ER-mitochondria (21.6±10.2 versus 35.4±9.3) interactions in HeLa *EXT1 k*.*d*. compared to control cells (Figures 3G-J). The latter was unexpected given the ∼5.7-fold increase in ER length (Figure 3B). However, it was found to correlate with an impaired calcium flux in those cells (Figure S3F-G), suggesting that cells undergo a metabolic switch following *EXT1 k*.*d*.

### EXT1 reduction induces Golgi re-organization and a metabolic switch

Although EXT1 is an ER-resident protein, it is also found in the Golgi apparatus, where it forms a catalytic heterodimer enzyme that polymerizes the elongation of heparan sulfate (HS) chains by sequential addition of glucuronic acid (GlcA) and *N*-acetylglucosamine (Glc-NAc) (Lind et al., 1998; McCormick et al., 2000; Senay et al., 2000). In addition to the notable changes in ER structure, TEM ultrastructural examination of *EXT1 k*.*d*. cells revealed structural changes in the Golgi apparatus size and shape (Figures 4A-B). The number of Golgi cisternae per stack was reduced (3.0±0.9 compared to 3.8±1.0 control; Figure 4C, Table S1), and stacks were dilated as well as shorter in length (729.2±329.0 compared to 1036.0±312.0 nm, respectively, Figures 4B and 4D, Table S1). Modified Golgi morphology combined with the reduction in ER-mitochondria interactions, points towards global metabolic changes in *EXT1 k*.*d*. cells.

**Figure 4.**
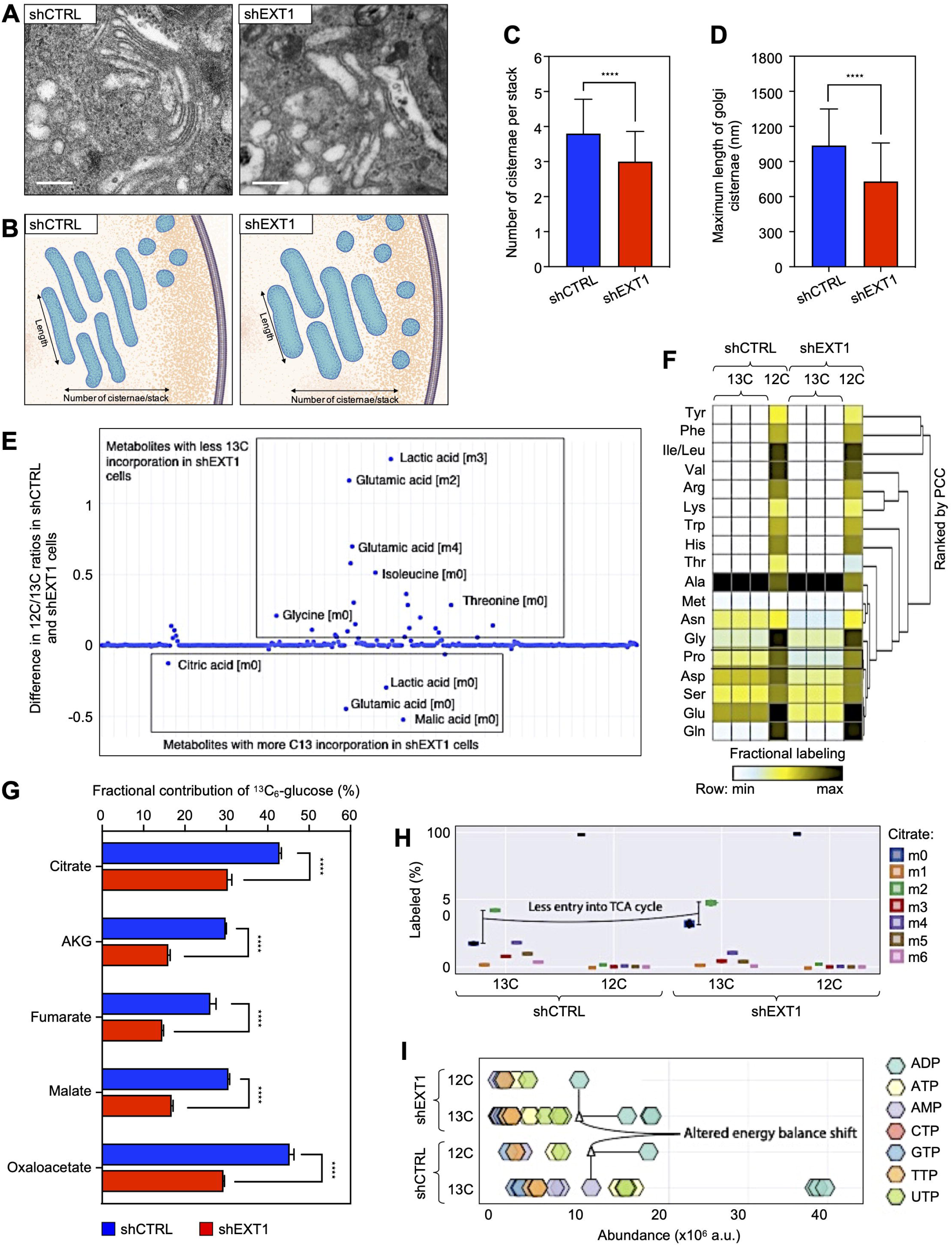
EXT1 reduction induces Golgi re-organization and a metabolic switch. **(A)** TEM of Golgi apparatus in HeLa shCTRL and shEXT1 cells. Silencing EXT1 results in reduced number, and increased size of Golgi cisternae/stacks. Scale bar, 500 nm. **(B)** Schematic representation of the Golgi apparatus as used for the statistical analysis of the different parameters. **(C-D)** (C) The number of Golgi cisternae/stacks and (D) and maximum length of individual Golgi cisternae (nm) was quantified based on TEM images (n = 20; Table S1). One-way ANOVA: ****p<0.0001. **(E)** Scatter plot of raw cell abundance profile of metabolites. **(F)** Heatmap Z-score represents fractional labeling of metabolites. Metabolites were clustered using one-minus Spearman’s rank correlation. **(G)** Fractional contribution from ^13^C_6_-Glucose to TCA metabolites (n = 3). One-way ANOVA: ****p<0.0001. **(H)** Isotopomer distribution of citrate derivatives into the TCA cycle in HEK293 cells (n = 3). **(I)** Cell abundance of nucleotides showing altered energy balance shift. See also Figure S4 and Table S1.

To assess the implications of EXT1 in cellular metabolism, we used two different strategies. First, we generated transcriptomic data (GSE138030) from cells treated with *EXT1* siRNA and control cells to reconstruct two *in silico* flux balance analysis (FBA) models using Constraint-Based Reconstruction Analysis (COBRA) (Heirendt et al., 2019) tools and the human *RECON2* metabolic model (Thiele et al., 2013). We found respectively 34 and 39 reactions uniquely active in the *EXT1 k*.*d*. or control models when the production of biomass was optimized (Figures S4A). These reactions are involved in the tricarboxylic acid (TCA) cycle, glycerophospholipid metabolism, pyruvate, methane, and sphingolipid metabolism (Figure S4B). Second, we performed high throughput metabolomic analysis of the relative abundance and fractional contribution of intracellular metabolites from major metabolic pathways in living *EXT1 k*.*d*. compared to control cells. We did not observe significant changes in glycolysis between control and *EXT1 k*.*d*. cells. However, in agreement with our *in-silico* FBA, we found that several nucleotides, amino acids, and metabolites from the TCA cycle were dysregulated in *EXT1 k*.*d*. cells (Figures 4E-I and S4C-E).

The fractional contribution of glucose carbons into these pools of metabolites was also decreased in *EXT1 k*.*d*. cells (Figure 4G). For instance, the fractional contributions of citric acid (change 12.51%, *p*<0.001), a-ketoglutarate (change 13.87%, *p*<0.0001), fumarate (change 11.61%, p<0.001), malate (change 13.74%, *p*<0.0001) and oxaloacetate (change 15.97%, *p*<0.0001) were significantly reduced in *EXT1 k*.*d*. cells (Figure 4G). Isotopologue profile analysis of TCA intermediates suggested that mitochondria were in a less oxidative mode of action in *EXT1 k*.*d*. cells, as m03, m04, m05, and m06 of citric acid were much lower in abundance (Figures 4H and S4E). In contrast, metabolite pools of the pentose phosphate pathway (PPP), the m05 of different nucleotides (ATP, UTP, GTP, and CTP), and the energy charge were increased in the *EXT1 k*.*d*. cells (Figures 4I and S4F-G). Altogether, these findings indicate a higher *de novo* synthesis and consumption rate of nucleotides necessary for the synthesis of sugar intermediates used in protein glycosylation (i.e., UDP-GlcNAc) in the *EXT1 k*.*d*. cells.

### *EXT1 k*.*d*. causes changes in the molecular composition of ER membranes

To understand the molecular mechanisms of the EXT1-mediated cellular metabolic changes observed above, we isolated ER microsomes from *EXT1 k*.*d*. and control cells. TEM revealed that, in the absence of EXT1, the structure of ER membranes was modified, as vesicle-like fragments were observed compared to the normal heterogeneous microsomes in control cells (Figure 5A).

**Figure 5.**
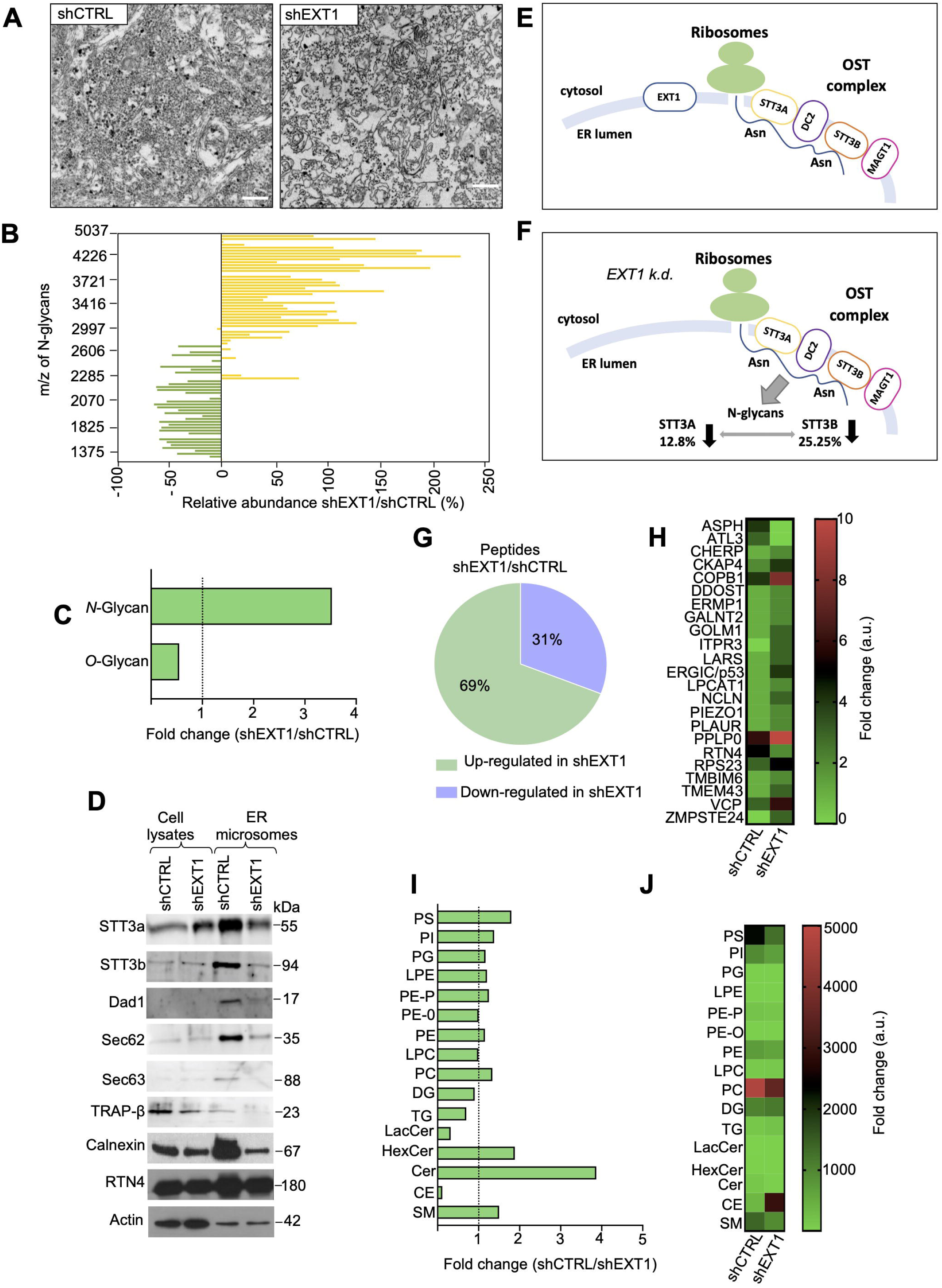
*EXT1 k*.*d*. causes changes in the molecular composition of ER membranes. **(A)** TEM of ER microsomes isolated from HeLa cells. Scale bar, 1 μm. **(B)** Glycomics analysis of microsomes. Relative abundance of each *N*-glycan in shEXT1 *vs*. shCTRL microsomes. The variations are plotted by *N*-glycan mass. One-way ANOVA: ***p<0.001, n.s., not significant. **(C)** As in (B), the bars indicate the fold change of the total *N*- and *O*-glycan intensities. **(D)** Expression of OST complex subunits (STT3a, STT3b, Dad1), other translocon members (Sec62, Sec63, Trap-β), and ER constitutive markers Calnexin and RTN4 in microsomes. **(E)** Schematic representation of the OST complex for which catalytic subunits STT3A and STT3B are less glycosylated following *EXT1 k*.*d*. **(F)** Quantitative proteomic analysis of microsomes. Pie chart illustrates the number of up-regulated and down-regulated proteins. **(G)** Heatmap to quantify 23 ER integral proteins. **(H)** Lipidomic analysis of different lipid species as found in microsomes. Bars indicate the fold change of the total intensity (a.u.). **(I)** As in (H) alternatively graphed in a heatmap. See also Figure S5, and Tables S2,S3, S4 and S5.

Based on this observation, we comprehensively compared the glycome, proteome, and lipidome profiles of those ER membranes in control and *EXT1 k*.*d*. cells. Glycome analysis using MALDI-TOF-MS enabled absolute and relative quantification of glycoprotein *N*- and *O*-glycan abundances, respectively in these cell lines (Table S2). The knockdown of *EXT1* did not induce the appearance of new glycan species on membrane proteins (Figure S5A). However, we observed a significant shift towards higher molecular weight *N*-glycans and more *O*-glycosylation compared to control ER membranes (Figures 5B-C, and Table S2). The total relative amount of *N*-glycans was reduced, consistent with the lower cellular abundance of UDP-GlcNAc (Figures 5C and S5B). This deregulation appears to occur, at least in part, at the level of the first step during protein *N-*glycosylation, which involves the OST complex. Indeed, microsomes isolated from *EXT1 k*.*d*. cells had reduced amounts of OST complex proteins STT3A, STT3B, and Dad1 (Figure 5D).

*N*-glycosylation in eukaryotes is co-translational (Kornfeld and Kornfeld, 1985; Bai et al., 2018; Braunger et al., 2018; Wild et al., 2018). In agreement with the finding that *EXT1 k*.*d*. impairs *N*-glycosylation, we observed that some members of the translocon complex (Sec62 and Sec63), the translocon-associated protein complex (TRAP), as well as the *N*-glycosylation quality control protein calnexin, were found to be reduced in *EXT1 k*.*d*. cells compared to controls (Figure 5D). Using a comparative mass spectrometry (MS/MS) analysis followed by glycopeptide identification, we found that *EXT1 k*.*d*. specifically reduced *N*-glycosylation of asparagine (N) residues N548 and N627 in STT3A and STT3B respectively, which are the catalytic subunits of the OST complex (Figures 5E-F, Table S3). Although the role of *N*-glycosylation of human STT3A and STT3B is still unknown, in yeast, *N*-glycosylation of the ortholog Stt3 mediates the assembly of the OST subcomplexes *via* interaction with Wbp1 and Swp1 (Bai et al., 2018).

MS/MS proteomic analysis also identified 226 proteins that were differentially abundant in ER membranes of *EXT1 k*.*d*. cells (Table S4), including 23 ER-resident proteins. Specifically, RTN4 and ATL3, ER-shaping proteins were found in lower abundance in *EXT1 k*.*d*. cells (Figures 5G-H). However, valosin-containing protein (VCP), an ATPase involved in lipid recruitment during ER formation (Zhang et al., 1994) and a glycan-binding component of the ER-Golgi intermediate compartment that is involved in ER reorganization, ERGIC/p53, were in higher abundance (Gaudet et al., 2011; Mitrovic et al., 2008; Nichols et al., 1998) (Figure 5H). Consistent with the increase in *O*-glycans (Figure 5B), ER membranes from *EXT1 k*.*d*. cells were also found to have higher amounts of GalNAc transferase 2 (GALNT2) (Figure 5H and Table S4) and an overall higher glycosyltransferase activity in ER microsomes (Figure S5C). These results confirm that the ER proteome, including shaping proteins and ER enzymes, is deregulated following *EXT1 k*.*d*.

Examination of lipid classes in ER microsomes also highlighted significant changes following EXT1 depletion (Figures 5I-J and Table S5). The most significant increase was observed in cholesterol esters (CE), which were ∼9-fold higher in *EXT1 k*.*d*. membranes compared to control (Figure 5I). Changes were also observed in phospholipids (PL) such as phosphatidylcholine (PC), phosphatidylserine (PS) and sphingomyelin (SM) (Figures 5I-J and Table S5). We concluded that, in the absence of EXT1, ER membrane lipid composition is modified towards structural fluidity (Lange et al., 1999).

### EXT1 localizes in ER tubules and sheet matrices

The above results suggest a role of EXT1 in the maintenance of the ER structure. Previous studies have shown that EXT1 localizes predominantly to the ER (McCormick et al., 1998, 1999). However, whether EXT1 localizes in ER tubules or sheet matrices was not investigated because of the spatial limitations of optical microscopy.

To precisely characterize EXT1 localization in ER structures, we performed super-resolution imaging (SR) (Veettil et al., 2008) with EXT1 tagged with SYFP2 and mEmerald, two fluorophores with different photostability properties. Using two SR technologies, stimulated emission depletion (STED) and structured illumination microscopy (SIM) (Hell and Wichmann, 1994; Schermelleh et al., 2010), we found that EXT1 localized in dense sheets and peripheral ER tubules (Figures 6A-B). EXT1 largely co-localized with the ER luminal marker protein disulfide isomerase family A member 3 (PDIA3) (Figures 6C) and, to lesser extents, with lectin chaperone calnexin as well as Golgi marker GM130 (Figures S6A-C). EXT1 perfectly colocalized with ER-shaping proteins Lunapark1 (Lnp1), ATL1, as well as RTN4a in tubules and the ER three-way junctions (Figures 6D-F and S6C).

**Figure 6.**
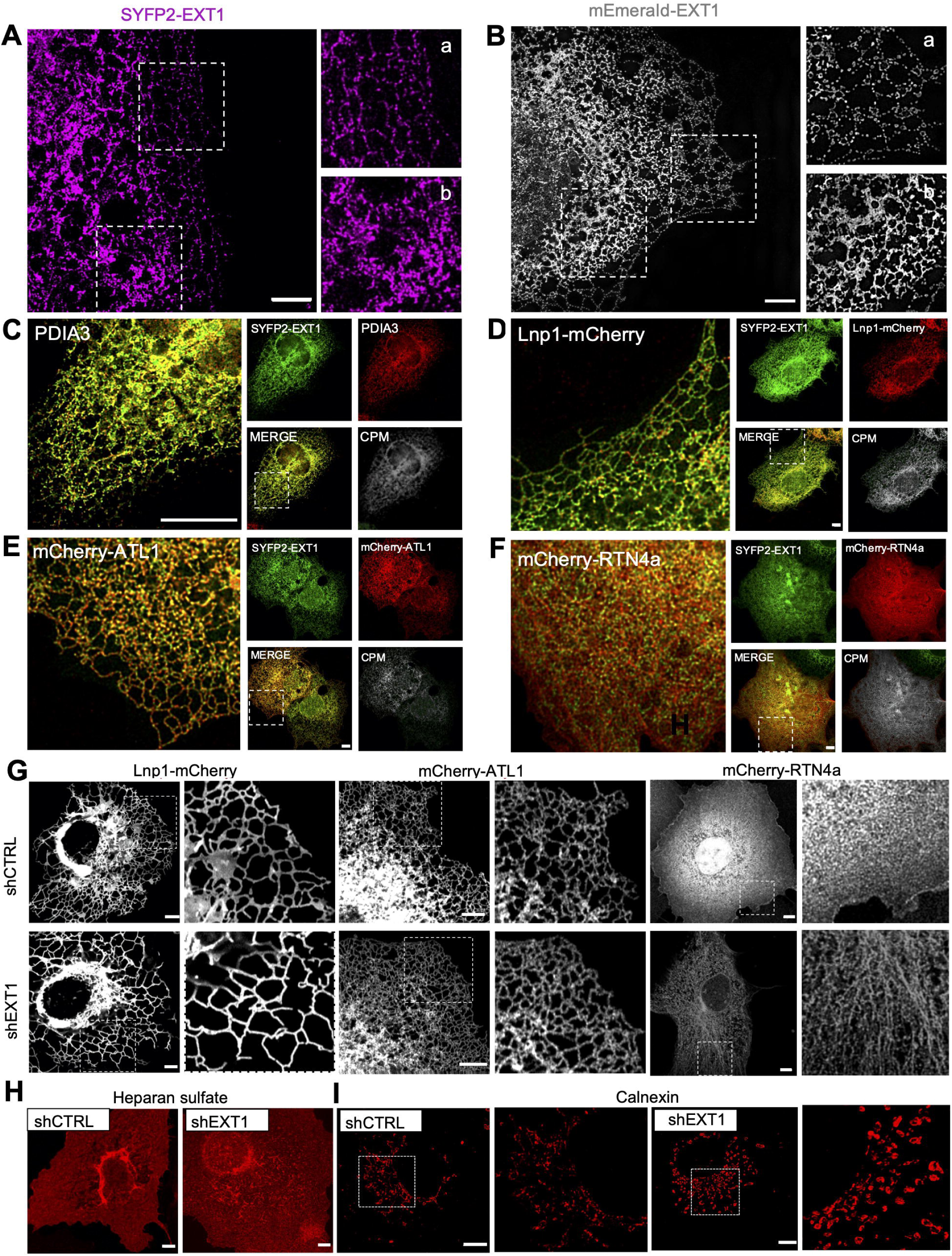
EXT1 localizes in ER tubules and sheet matrices. **(A-B)** STED (A) and SIM (B) images of Cos7 cells expressing SYFP2-EXT1 and mEmerald-EXT1, respectively. Boxed regions illustrate the tubular (subpanel a) and the cisternal (subpanel b) ER. Scale bar, 4 μm. **(C)** Confocal fluorescence microscopy of Cos7 cells transiently expressing SYFP2-EXT1 (green) and endogenous ER marker PDIA3 (red). The subpanels show the individual and merged channels and the Colocalized Pixel Map (CPM). Scale bar, 4 μm. **(D-F)** As in (C) but co-expression of SYFP2-EXT1 (green) and indicated ER markers (red). **(G)** Live imaging of shCTRL and shEXT1 Cos7 cells stably expressing indicated ER markers. Scale bar, 4 μm. **(H)** Heparan sulfate endogenous staining (red) of Cos7 cells. Scale bar, 5 μm. **(I)** Endogenous staining (red) of Cos7 cells with Calnexin antibody. Boxed regions magnified illustrate a zoom of circular, vesicle-like structures that appear following EXT1 knockdown. Scale bar, 4 μm. See also Figure S6.

To assess whether *EXT1 k*.*d*. might affect ER luminal dynamics, we analyzed the dynamic motion of ER tubules and three-way junctions by tracking the trajectories of ATL1 and Lnp1 proteins using live imaging (Video S1-4). In Cos7 *EXT1 k*.*d*. cells, the ER periphery morphology is asymmetrically dispersed compared to controls cells. While, Lnp1 and ATL1 markers showed increased ER tubular network, RTN4a showed increased membranous localization following EXT1 knockdown (Figures 6G). We quantified the ratio between ER tubules and three-way junctions, which indicated that the ER fusion rate was not affected following knockdown of EXT1 (Figure S6D). Next, we adapted a previously described single-molecule localization algorithm (Holcman et al., 2018) to reconstruct the diffusivity and velocities at the three-way junctions (Figures S6E-H). We computed particle distributions, trajectories, as well as velocities and found ATL1 to have a higher diffusivity and instantaneous velocity than Lnp1 (Figures S6G-H), consistent with their respective localizations in ER tubules and three-way junctions. The maximum tubular motion observed here (velocity of ∼3 µm/s) was lower than the luminal motion in previous observations (10-40 µm/s) (Holcman et al., 2018) (Figure S6H). EXT1 reduction by k.d. did not affect tubule motion, suggesting that the ER morphology changes in *EXT1 k*.*d*. cells might result from luminal flow changes, potentially driven by intracellular redistribution of heparan sulfate (Figure 6H).

The molecular chaperone calnexin, which assists protein folding in the ER, exhibited an aggregation pattern in *EXT1 k*.*d*. cells (Figure 6I), which might result in decreased movement of molecules through the ER lumen. To assess how a reduced polygonal area following *EXT1 k*.*d*. might influence ER luminal protein mobility and network continuity, we quantified the relative diffusion and active transport through the lumen of a photoactivable ER lumen marker (Jones et al., 2009) (PA-GFP-KDEL). Its signal was spread throughout the entire ER network, demonstrating that the continuity of ER was not affected in *EXT1 k*.*d*. cells (Video S5 and S6). However, we observed a significantly higher dynamic of fluorescence intensity in regions close to the nucleus (Figures S6I-J, at 8, 12, 16 µm), suggesting that the structural rearrangements of the ER following *EXT1 k*.*d*. actively participate in luminal protein transport. Altogether, these data demonstrate that *EXT1 k*.*d*. induces ER morphological changes that impair protein movement through the ER.

### *EXT1 k*.*d*. results in increased secretory cargo trafficking

To comprehensively assess the function of EXT1 in protein dynamics through the ER, we combined interactome analysis with imaging approaches. First, we captured the EXT1 interactome in ER microsomes by affinity purification and mass spectrometry analysis (Table S6). Consistent with a role in ER morphology, spatial analysis of functional enrichment (SAFE) (Baryshnikova, 2016) was used to identify three functional modules within EXT1 interactors, two of which were translation initiation and protein targeting to the ER (Figure 7A). Next, we investigated potential connections between EXT1 and the secretory pathway by comparing the proteome isolated from control and *EXT1 k*.*d*. cells after stable isotope labeling by amino acids (SILAC) (Figure S7B-C and Table S7). Differential protein expression analysis indicated the up-regulation of COPII anterograde vesicle-mediated transport components, with concomitant down-regulation of retrograde components following depletion of EXT1 (Figures S7D-E). Thus, we hypothesized that the depletion of EXT1 led to the enhancement of protein recruitment into luminal ER and subsequent secretion or distribution into cellular membranes.

**Figure 7.**
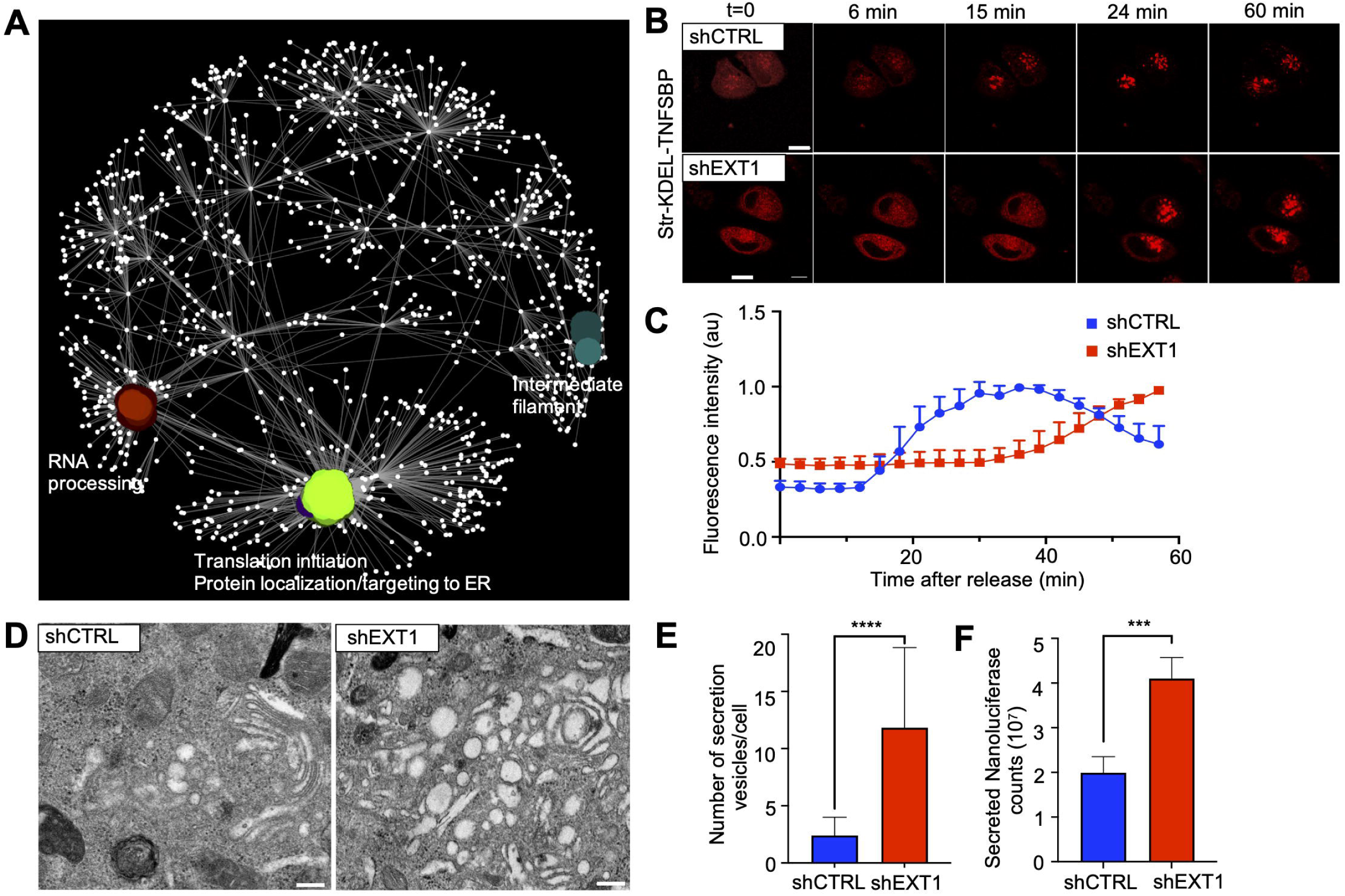
*EXT1 k*.*d*. results in increased secretory cargo trafficking. **(A)** SAFE analysis of EXT1 interactome in ER microsomes. **(B)** Live imaging of RUSH-synchronized traffic of TNF protein in HeLa cells. Scale bar, 10 μm. **(C)** Mean normalized fluorescence intensity (a.u.) after the addition of biotin. Two-stage linear step-up procedure of Benjamini, Krieger and Yekutieli. **(D)** TEM of *trans*-Golgi area of HeLa cells. Higher magnification of the boxed area is shown. Scale bar, 1 μm. **(E)** Number of secretion vesicles in the *trans*-Golgi area quantified based on TEM images (n = 17-18). Mean number + SD. One-way ANOVA: ****p<0.0001. **(F)**. Quantification of Nano-luciferase enzymatic activity from supernatant of shEXT1 and shCTRL Hela cells following transduction with a lentivirus delivering secreted Nanoluciferase. See also Figure S7 and Tables S6 and S7.

To further assess the changes in the secretory pathway, we monitored anterograde transport using the retention selective hook (RUSH) system (Boncompain et al., 2012) that enables the synchronization of cargo trafficking. By tracking cargo transport from the ER to the Golgi using live imaging, we observed a slower dynamic response in *EXT1 k*.*d*. cells that resulted in an increased residency of the cargo within the secretory pathway (Figures 7B-C and Video S7 and S8). This finding was confirmed using an additional ER export assay based on the vesicular-stomatitis-virus glycoprotein (VSVG) (Wilhelmi et al., 2016) (Figures S8A-B), and by examining COPII coat structural components SEC16 and SEC31 (Figures S7C-D). TEM analysis also indicated a higher number of *trans*-Golgi secretory vesicles (11.83±7 and 2.4±1.6 secretion vesicles/cell, in *EXT1 k*.*d*. and control cells, respectively) following depletion of EXT1 in HeLa cells (Figures 7D-E). Finally, we confirmed enhanced secretion by producing significantly more recombinant proteins (Nano-luciferase) in HeLa *EXT1 k*.*d*. compared to control cells (Figure 7F). The integration of all the above results allowed as to conclude that EXT1 controls secretion by interacting with components of the general translational initiation machinery (Figure S7A). Thus, EXT1 expression reduction, as modeled by SAFE analysis combining quantitative transcriptional and translational expression (Figure S8E-F), affects several processes in mammalian cell physiology and metabolism.

## DISCUSSION

Multiple pieces of evidence indicate that EXT1 is broadly implicated in cancer, as suggested by the findings shared in the Cancer Cell Line Encyclopedia (CCLE) (Ghandi et al., 2019), cancer genome atlas (TCGA) (Lorio et al., 2016) and catalogue of somatic mutations in cancer (COSMIC) (Tate et al., 2018; McDonald et al., 2017). *EXT1* mutations range from 1% in small cell lung cancer (SCLC) tumors to 27% in colorectal cancers (COREAD) (Figure S1). At the protein level, we previously reported that EXT1 interacts with Notch1 (Daakour et al., 2016). Here we provide data showing that *EXT1* gene should be considered as an additional regulator molecule involved in T lymphocytes development in mice (Figures 1B-E). Furthermore, our results from *EXT1* and *Notch1* double k.o. in developing thymocytes, demonstrate that cells are able to pass the critical β and γδ-selection checkpoints in the absence of *Notch1* expression (Figure 1 D-E). This genetic suppression interaction between *EXT1* and *Notch1* in developing thymocytes reflect a mechanism (exocytosis) potentially controlling T-lineage specification, in addition to the well-known transcription and ligand-receptor modulations (Radtke et al., 2013). Genetic suppression is one of the most powerful tools in yeast (Van Leeuwen et al., 2016) and *C. elegans* (Wu and Han, 1994; Zheng et al., 2004) genetics. In these organisms, genetic suppression is facilitated by the ability to generate and handle a large number of individual mutations *in vivo*, allowing global scale connection of genes involved in the same pathway or biological process. Although systematic examination of *EXT1* genetic interactions was impractical in our mouse models, we demonstrated an overlapping role of *EXT1* and *Notch1* in the developmental stages (DN3 and DN4) of thymocytes, and an unexpected healthy phenotype of thymocytes with *EXT1*^*F/F*^ *Notch1*^*F/F*^ double knockout. We thus validated a physiological genetic suppression role of *EXT1* in the function of the Notch1 transmembrane receptor. To further demonstrate a potential global role of EXT1 in cancer, we took advantage of the synthetic lethality (SL) (Feng et al., 2019; Jerby-Arnon et al., 2014; Lee et al., 2018a), and the synthetic dosage lethality (SDL) (Megchelenbrink et al., 2015) principles, whereby for each pair, individual gene inactivation (SL) or expression variation (SDL) result in viable phenotypes, whereas combined perturbations are lethal. These approaches identified several oncogenes, including *KRAS, CREBBP, PTEN*, and *BRCA2*, as SDL genetic partners of *EXT1*, highlighting its potential clinical relevance in different cancers.

*In vitro*, the formation of the ER tubular network requires only a small set of membrane-curvature and stabilizing proteins (RTNs, REEPs, and large ATL-GTPases) (Powers et al., 2017). However, these effectors cannot account for the diversity and adaptability of ER size and morphology observed in individual cell types. It is expected that *in vivo*, the dynamics of tubular three-way junctions and tubule rearrangements to accommodate luminal flow mobility, rely on additional proteins or mechanisms (Chen et al., 2012). Despite the discovery of glycoproteins in intracellular compartments 30 years ago (Kelly and Hart, 1989), our knowledge about the glycoproteome is still biased towards secreted and plasma membrane proteins. Glycosylation is well known to regulate the physical properties of different glycolipid and glycoprotein biopolymers at the surface of mammalian cells by controlling plasma membrane and cell coat morphologies (Shurer et al., 2019). The central enzyme in the *N*-glycosylation pathway is the oligosaccharyltransferase (OST) complex, which catalyzes the transfer of oligosaccharides from dolichol pyrophosphate-linked oligosaccharide to the nascent polypeptides in the protein translocon systems (Kelleher and Gilmore, 2006; Schnell and Hebert, 2003). The spatial organization of these protein modules is tightly regulated to coordinate temporally coupled synthesis, *N*-glycosylation, and protein translocation (Braunger et al., 2018; Wild et al., 2018). The atomic structure of yeast OST complex highlighted a potential role of an *N*-glycan at the N539 position of the catalytic subunit STT3 in the stability of the OST complex by sticking together Wbp1 and Swp1 interacting subunits (Bai et al., 2018). Here, we observed that glycosylation of the corresponding residues N548 and N627 of STT3A and STT3B mammalian OST is impaired following the reduction of EXT1. This suggests that EXT1 is involved in the stability of the OST complex in the ER lumen, providing a mechanistic explanation for the lower *N*-glycome observed in *EXT1 k*.*d*. cells (Figure 5C). The resulting alternative glycosylation pattern of ER membrane proteins observed here correlated with extensive ER architectural and functional remodeling. Our results suggest that EXT1 is topologically involved in the stability of the OST and the translocon complexes in the ER.

The results presented here provide insights into a specific fundamental downstream role of EXT1 in the architecture of the ER. We demonstrated that the reduction of EXT1 affects ER structures, membrane glycome, and lipid compositions, which have broad-ranging metabolic consequences for the cell. EXT1 expression reduction, which not only affects the structure of the ER, also favors membrane structural fluidity and affects its luminal dynamics. Thus, our findings have demonstrated that glycosylation is an important post-translational modification controlling the internal plasticity and structure-function of the ER. In the future, it will be interesting to identify other glycosyltransferases that co-regulate intracellular organelle morphologies. It is also essential to determine the atomic structure of EXT1 at the ER, in order to clarify the positioning of EXT1 relative to the OST and translocons complexes. Taken together, our findings suggest that the diversity of proteoglycans destined to the cell surface results from the glycosylation equilibrium of intracellular and plasma membrane proteins. At the fundamental level, our findings argue for a general biophysical model of ER membrane-extension and functions regulated by resident glycosyltransferase enzymes like EXT1.

## ACKNOWLEDGMENTS

We are grateful to Yu Yamaguchi (Sanford Children’s Health Research Center Sanford-Burnham Medical Research Institute, La Jolla, CA, USA) and Freddy Radtke (Ecole Polytechnique Federale de Lausanne, Lausanne, Switzerland) for providing EXT1/loxP and NOTCH1-floxed mice, respectively. We thank the following investigators for providing essential plasmids: Tom Rapoport (Dept of Cell Biology, Harvard Medical School, MA, USA) for Atl1, Rtn4a, and Lnp mCherry fusion plasmids; Jennifer Lippincott-Schwartz (Janelia Research Campus, Howard Hughes Medical Institute (HHMI), Ashburn, VA, USA) for mEmerald-Sec61b; Florian Heyd (Department of Biology, Chemistry, Pharmacy, Freie Universitat Berlin, Germany) for VsVg construct; Vincent Timmerman (Peripheral Neuropathy Research Group University of Antwerp, Belgium) for Atl3-mCherry construct, Vicky C Jones (University of Central Lancashire, Preston, UK) for the LV-PA-KDEL-GFP construct, Richard Zimmermann (Medical Biochemistry and Molecular Biology, Saarland University, Homburg, Germany) for translocon antibodies. We also thank the Eurobioimaging nodes at Maastricht (Netherlands) and Turku (Finland) Universities for super-resolution STED and SIM imaging, respectively. We thank the following University of Liege core facilities: In vitro Imaging, Mouse facility and transgenics, Proteomics and Viral vectors for their services. We thank Patricia Piscicelli for technical assistance in TEM. We thank Jonas Dehairs and KU Leuven Departement of Oncology for Lipidomics analysis. D-K. K. was supported by Banting Postdoctoral Fellowship of Canada and Basic Science Research Program through the National Research Foundation of Korea (NRF) funded by the Ministry of Education (2017R1A6A3A03004385). B.S.D., S.D., D.R.N., A.J., D.C.J-E.A., and K.S-A. were supported by New York University Abu Dhabi (NYUAD) Institute grant 73 71210 CGSB9 and NYUAD Faculty Research Fund AD060. D.K. was supported by an FRS-FNRS-Télévie Fellowship #7651317F (J-C.T). J-C.T is Maitre de Recherche of the FRS-FNRS. Primarily the Fonds de la Recherche Scientifique (FRS-FNRS) and the Fonds Leon Fredericq grants supported this work.

## AUTHOR CONTRIBUTIONS

The project was conceived and supervised by J-C.T. and F.D. Major experiments were performed by D.K. Validation experiments were performed by D.K., M.T., F.M., P.L., M.H., J.C., B.G., and D.V. supervised by K.K., M.T., P.V.V., C.D., M.V.Z, J-C.L., C.L-I. and J-C.T. Image analysis was performed by D.K., C.P. and N. d-C supervised by V.K. and J-C.T. Downstream computational analyses were performed by B.S.D., D.R.N., S.D., D.S., and D-K.K. supervised by K.S-A and J-C.T. The paper was written by D.K., K.J.L. and J-C.T. with help from J.O, J.H., D.N., K.S-A., M.V. and F.D.

## SUPPLEMENTAL FIGURE LEGENDS

**Figure S1. *EXT1* is a suppressor hub in different cancer types, Related to Figures 1 and 2**

**(A-B)** Relative mRNA expression levels of mouse *EXT1* (A) and mouse *Notch1* (B) in thymocytes isolated from indicated mice. One-way ANOVA: ****p<0.0001; ns: not significant. **(C-D)** Western blot analysis of intracellular Notch1 (ICN1) in Jurkat-luciferase shEXT1 (C) or Jurkat-luciferase EXT1-GFP (D) compared to control cells. **(E)** Proliferation of Jurkat cells measured with BrdU optical absorbance at 450 nm. Mean number + SD is plotted. One-way ANOVA: ns: not significant. **(F)** Network depicting the synthetic lethality (SL) interactome map of EXT1. Blue edges indicate SL observed experimentally and verified clinically, while red edges indicate SL not verified clinically. **(G-H)** Pie chart representing the percentage of EXT1 mutations in cancer cell lines (G) and in tumor samples (H). Data are from TCGA and COSMIC databases.

**Figure S2. EXT1 is the only member of the EXT family regulating ER structures in different cell lines, Related to Figure 2**

**(A)** TEM of ER morphology in HEK293 cells. Scale bar, 2 μm. **(B)** The relative mRNA expression level of *EXT1* gene was analyzed by qPCR in HEK293 cells. One-way ANOVA: ****p<0.0001. **(C)** As in (A) for Jurkat cells. **(D)** As in (B) for Jurkat cells. **(E)** TEM of ER structure of HeLa cells depleted for *EXT1, EXT2, EXTL1, EXTL2*, or *EXTL3* expression. Scale bar, 2 μm. **(F-J)** Relative mRNA expression levels of the *EXT1* (F), *EXT2* (G), *EXT-L1* (H),*-L2* (I), *and -L3* (J) genes analyzed by qPCR. One-way ANOVA: *p<0.05, **p<0.01, ***p<0.001.

**Figure S3. EXT1 reduction induces ER tubules re-organization and a shift in calcium flux, Related to Figures 2 and 3**

**(A)** Proliferation of HeLa cells measured with BrdU optical absorbance at 450 nm. Mean number + SD is plotted. One-way ANOVA: ns: not significant. **(B)** Xcelligence system to compare cell indexes. **(C-E)** Quantitative analysis based on the skeletonization model of Cos7 cells expressing Sec61b. (C) Tubule mean length. Box plot indicates the mean and whiskers show the minimum and maximum values (n = 19-24). (D) As in (C) but for the cisternal mean area. (E) As in (C) but for the perimeter mean length. **(F)** Fluo-4 dye was used to measure Ca2+ concentration in Cos7 shCTRL and shEXT1 cells. **(G)** Mean normalized fluorescence intensity (a.u.) was measured (n = 12). Boxplot indicates the mean and whiskers show the minimum and maximum values. One-way ANOVA: ***p<0.001.

**Figure S4. EXT1 reduction induces a metabolic switch, Related to Figure 4**

**(A)** Venn diagram showing active reactions in control model (blue) and *EXT1* knocked-down model (red). **(B)** Pathways enriched in the active reactions. Blue, Red and black correspond respectively, to reactions uniquely in shCTRL, shEXT1, and in both models. **(C)** Metabolomic analysis from ^13^C_6_-Glucose of glycolysis nucleotide metabolites in HEK293 cells. Fold change in the abundance of the metabolites in shEXT1/shCTRL is shown. **(D)** As in (C) for amino acid metabolites. **(E)** As in (C) for citrate metabolites. **(F)** Percentage of energy charge. Energy Charge is calculated as (ATP+(0.5*ADP))/SUM(ATP+ADP+AMP), (n = 3). **(G)** As in (C) for pentose phosphate pathway metabolites. One-way ANOVA: *p<0.05, **p<0.01, ***p<0.001, ****p<0.0001, n.s., not significant.

**Figure S5. *EXT1 k*.*d*. causes changes in the glycome composition and glycosylation of ER membranes, Related to Figure 5**

**(A)** *N*-glycans profiles of microsomes isolated from HeLa cells shCTRL and shEXT1.

**(B)** Cell abundance from ^13^C_6_-Glucose of UDP-GlcNAc from ^13^C_6_-Glucose in HEK293 cells. One-way ANOVA: ***p<0.001, ****p<0.0001, n.s., not significant. **(C)** Phosphatase-coupled glycosyltransferase assay was used to compare the glycosyltransferase activities from microsomes. In the bar graph the mean number + SD is plotted. One-way ANOVA: ***p<0.001, n.s., not significant.

**Figure S6. EXT1 colocalizes with luminal and shaping ER markers, and its depletion does not affect luminal diffusivity, Related to Figures 6**

**(A)** Colocalization between EXT1 (green) and Calnexin (red). The subpanels show the individual and merged channels and the Colocalized Pixel Map (CPM). Scale bar, 4 μm. **(B)** As in (A) but GM130 (red). **(C)** Average Pearson’s correlation coefficient of indicated markers and EXT1 protein. **(D)** The ratio of the tubules over junctions in cells expressing ATL1 or Lnp1. **(E)** Diffusion analysis flowchart. **(F)** The average diffusivity in cells expressing ATL1 and Lnp1. **(G)** Flowchart of image processing for velocity measurements. **(H)** Analysis of velocities of the junctions in cells expressing ATL1 and Lnp1. **(I)** Live imaging of activated PA-KDEL-GFP. Scale bar, 5 μm. **(J)** Ribbon plots show the mean fluorescence intensity ± SEM at fixed distances (8, 12 and 16 μm) from the photoactivation site. Two-way ANOVA followed by Sidak’s multiple comparisons test.

**Figure S7. EXT1 depletion is associated with increased translation and secretion, Related to Figure 7**

**(A)** Efficient depletion of EXT1 by shRNA. **(B)** Schematic representation of the SILAC workflow. **(C-D)** Up (C) and down-regulated (D) proteins involved in anterograde and retrograde transport. **(E)** The localization of a ts045-VSVG-GFP reporter during different time points. Beta-catenin (red) is used for plasma membrane labeling, scale bar, 5 μm. **(F)** Quantification of secretion. Mean number + SD. One-way ANOVA: **p<0.01, ***p<0.001, n.s., not significant. **(G)** Cos7 cells expressing indicated COPII coat markers. Scale bar, 4 μm. **(H)** As in (G) for SEC31A. Scale bar, 4 μm. **(I)** SAFE analysis combining de-regulated genes at transcriptional level, and differential protein abundance. Functional modules identified after enrichment (left). Merge of the two highlighting enriched functional modules (right).

## METHODS

Detailed methods of this paper and include the following:

- KEY RESOURCES TABLE
- LEAD CONTACT AND MATERIALS AVAILABILITY
- EXPERIMENTAL MODEL AND SUBJECT DETAILS
  ○ Mice
  ○ Cell lines

- METHODS DETAILS
  ○ Mice generation
  ○ Mice cell preparation, in vitro T-cell activation and polarization
  ○ RNA extraction and RT-qPCR
  ○ BrdU proliferation assay
  ○ Proliferation assay using the xCelligence system
  ○ Subcutaneous xenograft studies
  ○ Transmission electron microscopy
  ○ Flow cytometry, extracellular and intracellular staining
  ○ Plasmids
  ○ Mammalian cell lines generation and culture
  ○ DNA-siRNA transfection
  ○ Calcium Flux Detection assay
  ○ Preparation of microsomes from cultured cells
  ○ Western blotting and antibodies
  ○ N-glycans and O-glycans profiling
  ○ Metabolomics profiling
  ○ Lipidomics
  ○ Glycosyltransferase assay
  ○ Immunofluorescence and confocal, super-resolution microscopy
  ○ Photoactivatable GFP Imaging
  ○ Affinity purification for mass spectrometry
  ○ Mass spectrometry
  ○ SILAC labeling
  ○ Rush assay
  ○ Export assay

- QUANTIFICATION AND STATISTICAL ANALYSIS
  ○ Image analysis
  ○ SAFE analysis
  ○ Model generation and flux balance analysis
  ○ Statistical analysis

- DATA AVAILABILITY

## KEY RESOURCES TABLE

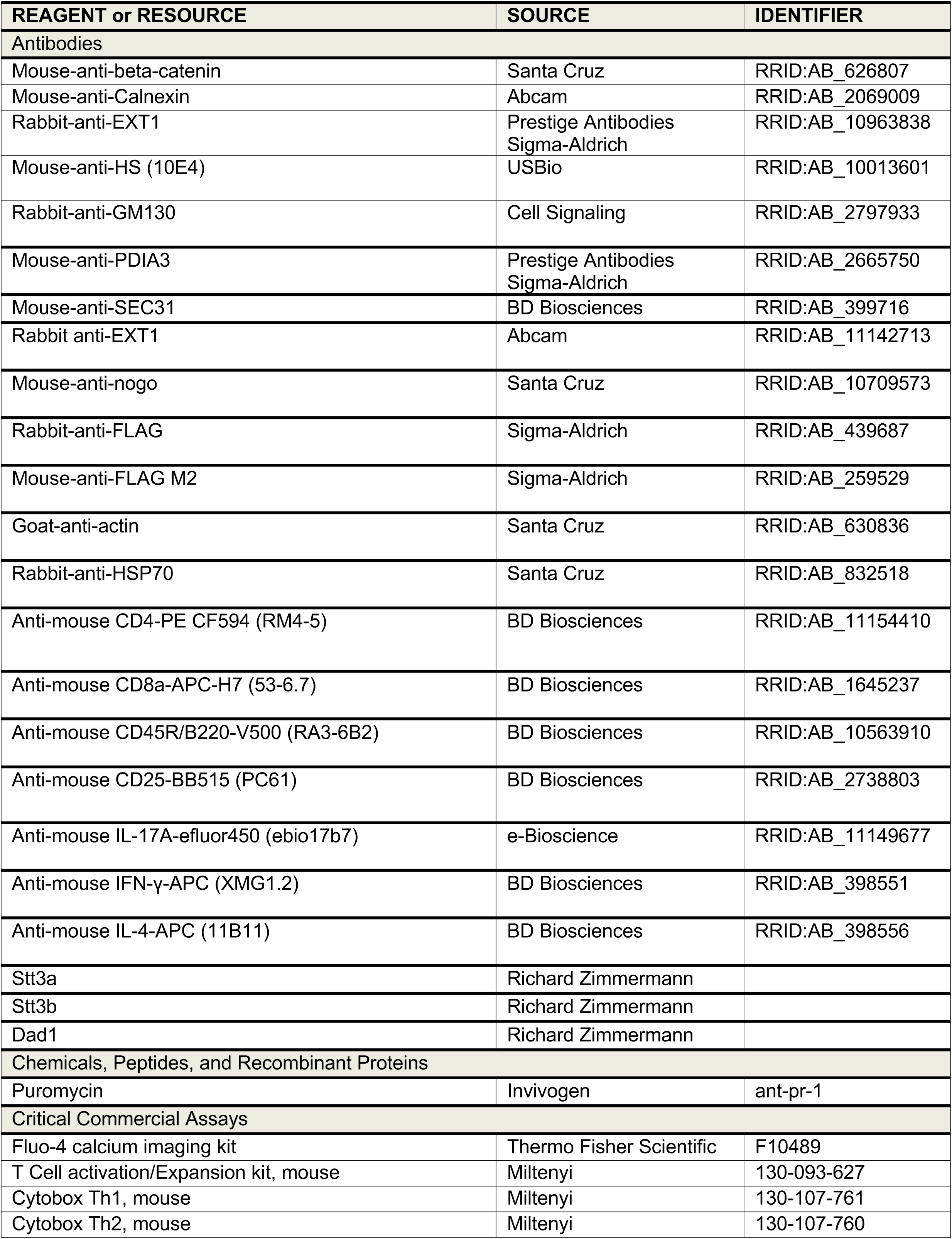
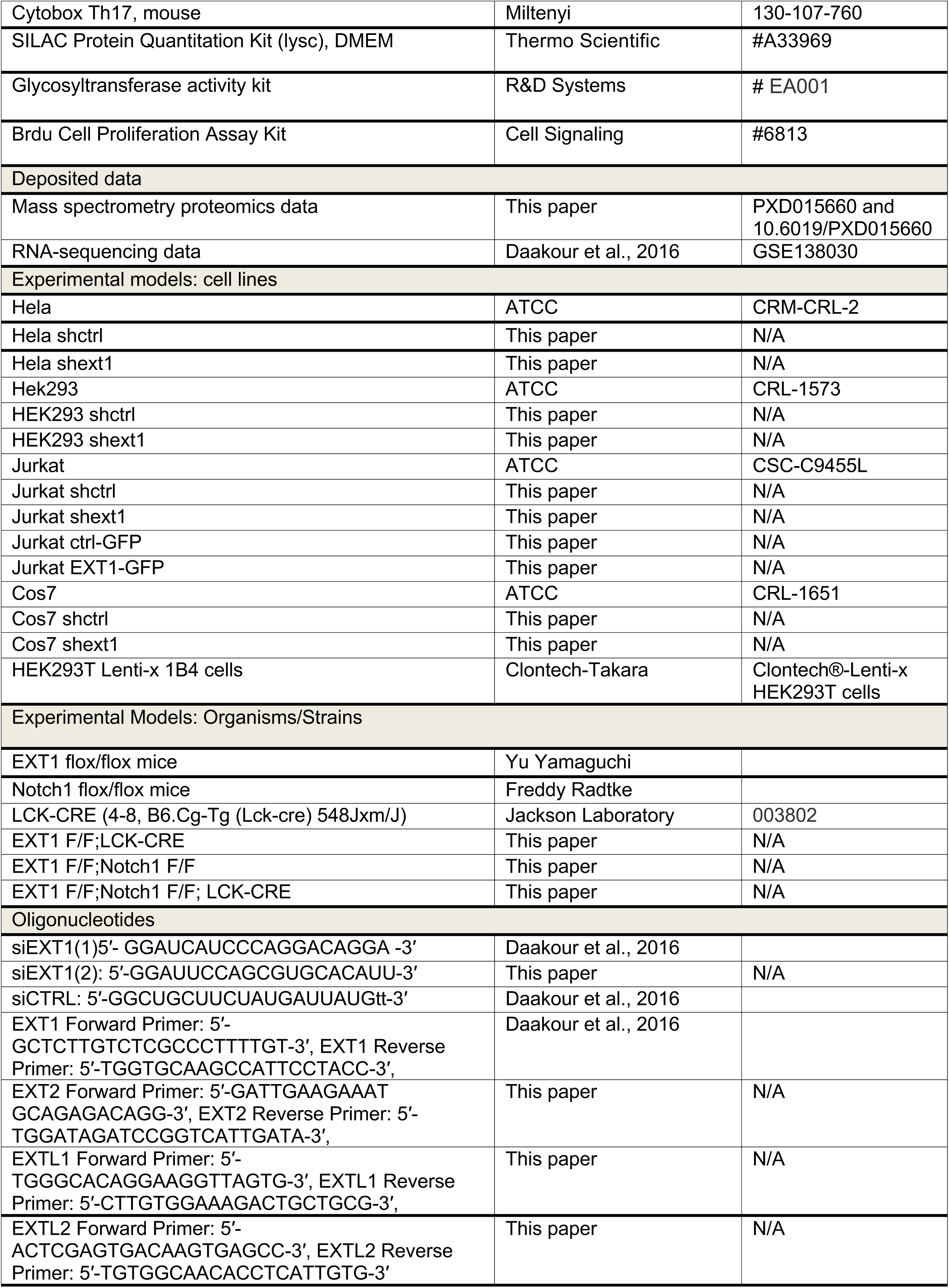
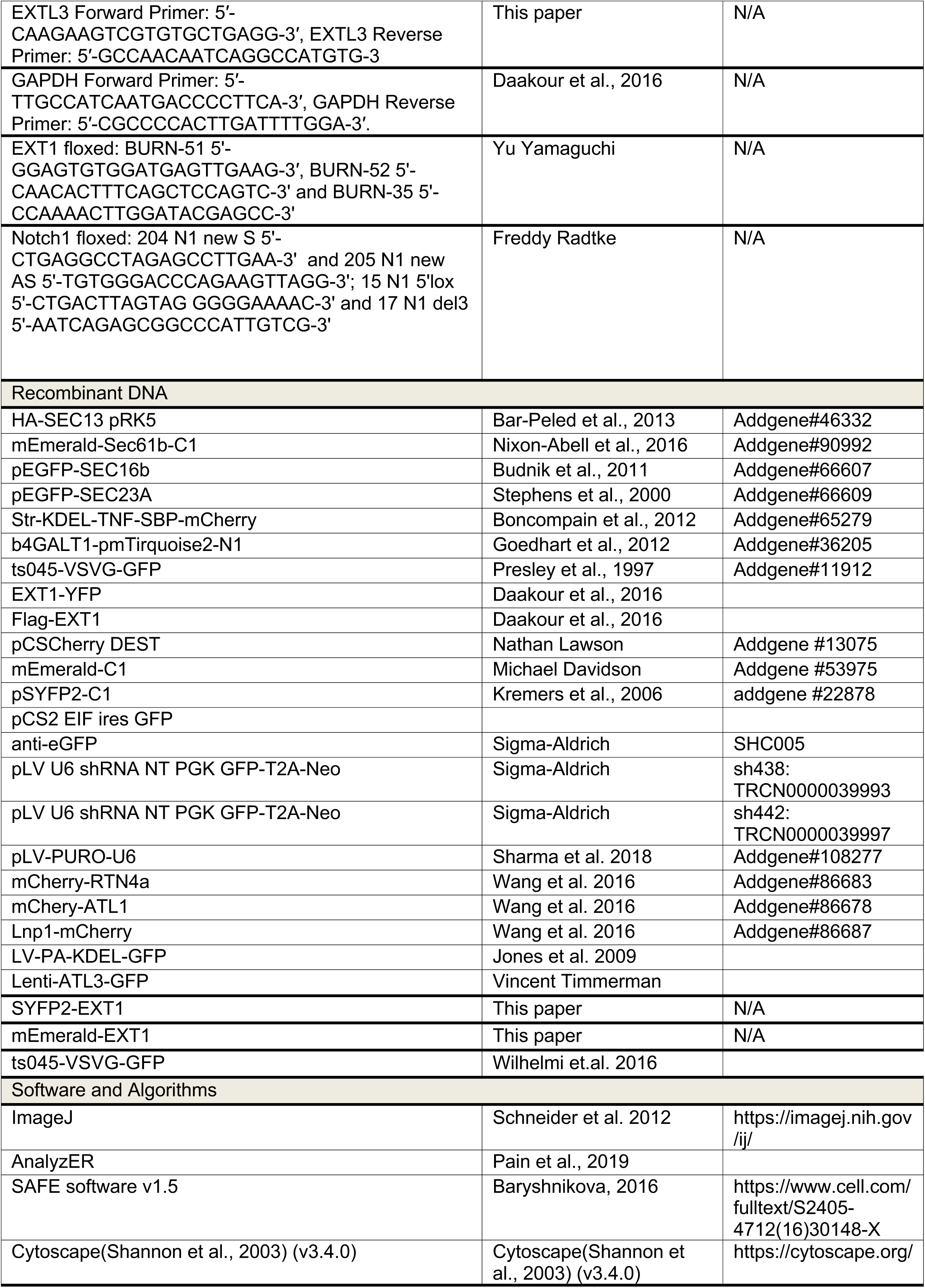
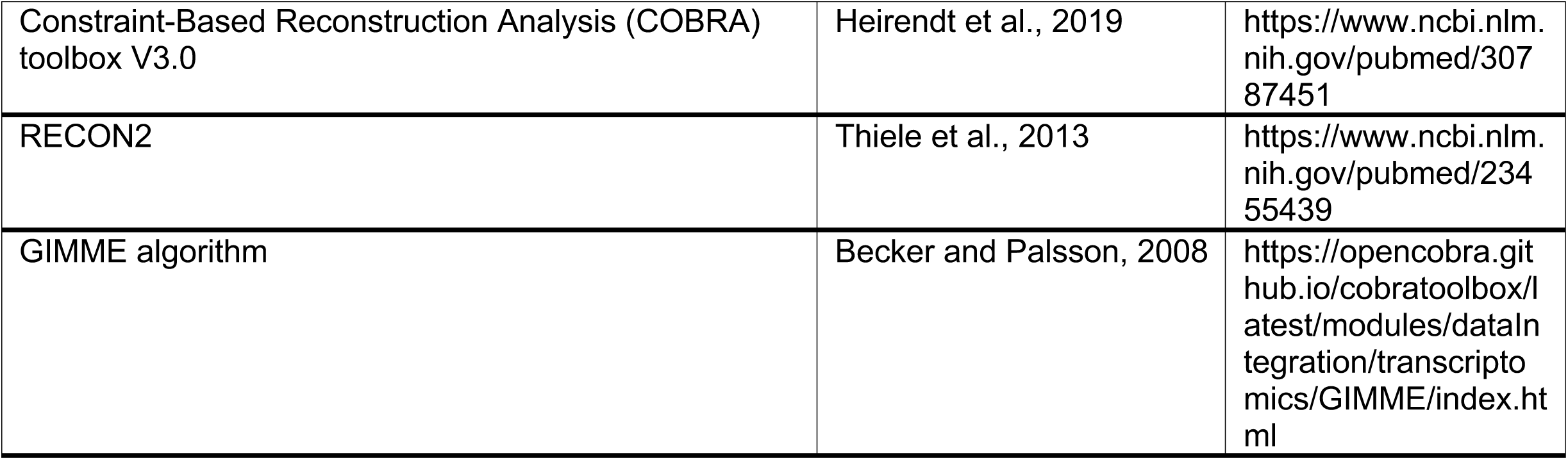

### CONTACT FOR REAGENT AND RESOURCE SHARING

Further information and requests for resources and reagents should be directed to and will be fulfilled by the Lead Contact, Dr. Jean-Claude Twizere. Plasmids and cell lines generated in this study are available upon request and approval of the Material Transfer Agreement (MTA) by the University of Liege.

### EXPERIMENTAL MODEL AND SUBJECT DETAILS

#### Mice

C57BL/6J background mice *EXT1*^*lox/lox*^, *Notch1*^*lox/lox*^, *lck-cre* and mice obtained after intercrossing were housed in the Animal Facility of the University of Liege. The protocol was approved by local ethical committee (authorization #13-1586).

#### Cell lines

HEK293 (*Homo sapiens*, fetal kidney), HeLa (Homo sapiens, cervical cancer) and Cos7 (African green monkey kidney) cells were cultured in Dulbecco’s Modified Eagle Medium (DMEM) supplemented with 10% fetal bovine serum, 2mmol/L L-glutamine and 100 I.U./mL penicillin and 100μg/mL streptomycin. Cells were incubated at 37°C with 5% CO2 and 95% humidity. Jurkat (Homo sapiens, T-cell leukemia) cells were cultured in Roswell Park Memorial Institute (RPMI) supplemented with 10% fetal bovine serum, 2mmol/L L-glutamine and 100 I.U./mL penicillin and 100 μg/mL streptomycin. Cells were incubated at 37°C with 5% CO2 and 95% humidity.

### METHOD DETAILS

#### Mice generation

T-cell specific deletion of *EXT1, Notch1*, or both genes on a C57BL/6J background was accomplished by intercrossing the EXT1 flox allele (Inatani et al., 2003) or Notch1 allele (Radtke et al., 1999) and the *lck-cre* transgene (Lee et al., 2001). *EXT1* flox/flox and Notch1 flox/flox mice were a gift from Dr. Yu Yamaguchi (Sanford Children’s Health Research Center Sanford-Burnham Medical Research Institute, La Jolla, CA, USA) and Freddy Radtke (Ecole Polytechnique Federale de Lausanne, Lausanne, Switzerland), respectively. LCK-CRE (4-8, B6.Cg-Tg (Lck-cre) 548Jxm/J) mice were purchased from Jackson Laboratory. T-cell specific depletion of *EXT1* and *Notch1* were verified by PCR. To detect deletion of the *EXT1* gene by PCR, the following primers were used on tail tips: BURN-51 5’-GGAGTGTGGATGAGTTGAAG-3′, BURN-52 5’-CAACACTTTCAGCTCCAGTC-3’ and BURN-35 5’-CCAAAACTTGGATACGAGCC-3’. BURN-51 and BURN-52 generate a 460 bp fragment from the wildtype allele and BURN-51 and BURN-35 a 509 bp fragment from CRE-excised allele. To detect Notch1 deletion by PCR, we used the following primers 204 N1 new S 5’-CTGAGGCCTAGAGCCTTGAA-3’, 205 N1 new AS 5’-TGTGGGACCCAGAAGTTAGG-3’; generating a 500 bp floxed fragment and a 450 bp in wild-type. We also used 15 N1 5’lox 5’-CTGACTTAGTAG GGGGAAAAC-3’ and 17 N1 del3 5’-AATCAGAGCGGCCCATTGTCG-3’ detecting a deleted band at 400bp. LCK-CRE mice were used as wild-type controls.

For RT-qPCR on thymocytes isolated from *EXT1*^*F/F*^*/lck-cre, Notch1*^*FF*^*/lck-cre, Notch1*^*F/F*^ *EXT1*^*F/F*^*/lck-cre* and *lck-cre* control mice, the following primers were used mEXT1, 5’-GCCCTTTTGTTTTATTTTGG-3’ and 5’-TCTTGCCTTTGTAGATGCTC-3’; mNotch1, 5’-GACACCTCTGGACAACGCCT-3’ and 5’-CGTGCTCACAAGGGTTGGCAC-3’; GAPDH, 5’-CCAGTATGACTCCACTCACG-3’ and 5’-GACTCCACGACATACTCAGC-3’. All experiments were done with mice in the C57BL/6 background. The protocol was approved by the University of Liege ethical committee (authorization #13-1586).

#### Mice cell preparation, *in vitro* T-cell activation and polarization

Single-cell suspensions from spleen, lymph node, blood, and thymus were obtained by mechanical disruption, straining over a 40 mm nylon mesh, and lysis of erythrocytes. Cells were counted and further stained as described in the section below, describing mice antibodies and flow cytometry. For primary T cell activation, polarization and cytokine detection of wild type or KO mice, isolated CD4+ T cells from spleen and lymph nodes by a negative magnetic separation (MACS) using CD4+ T cell isolation kit (Miltenyi Biotech) cultured at a density of 1×10^6^ cells per mL on 96-well plate for 72 h with biotinylated CD3 and CD28 antibodies (T Cell activation/Expansion kit, Miltenyi) in RPMI medium supplemented with 50 units/ml IL-2 (Miltenyi). Polarization to Th1, Th2, and Th17 was performed using CytoBox Th1, CytoBox Th2, and CytoBox Th17, respectively for 5 days. For Th1 population the medium was supplemented with cytokines and antibodies: 10 ng/mL (60 U/mL) mouse IL-12, 10 ng/mL (50 U/mL) mouse IL-2 and 10 µg/mL anti-IL-4 pure-functional grade. For Th2 population the medium was supplemented with the following cytokines and antibodies: 10 ng/mL (200 U/mL) mouse IL-4, 10 ng/mL (50 U/mL) mouse IL-2 and 10 µg/mL anti-IFN-γ pure-functional grade. For Th17 the medium was supplemented with 20 ng/mL (10000 U/mL) mouse IL-6, 10 ng/mL (40 U/mL) mouse IL-23, 10 ng/mL (8400 U/mL) mouse IL-1β, 2 ng/mL (10 U/mL) human TGF-β1, 10 µg/mL anti-IL-4 pure-functional grade, 10 µg/mL anti-IFN-γ pure-functional grade and 10 µg/mL anti-IL-2 pure-functional grade.

#### RNA extraction and RT-qPCR (human cell lines)

For expression studies, total RNA was extracted from the cell pellet using Nucleospin RNA kit (Macherey-Nagel) according to the manufacturer’s instructions. Real-time qPCR was performed using LightCycler® 480 SYBR Green I Master (Roche) and analyzed in triplicate on a LightCycler (Roche). The relative expression levels were calculated for each gene using the ΔΔCt method with GAPDH as an internal control. Primer sequences for qPCR are: EXT1 Forward Primer: 5′-GCTCTTGTCTCGCCCTTTTGT-3′, EXT1 Reverse Primer: 5′-TGGTGCAAGCCATTCCTACC-3′, EXT2 Forward Primer: 5′-GATTGAAGAAAT GCAGAGACAGG-3′, EXT2 Reverse Primer: 5′-TGGATAGATCCGGTCATTGATA-3′, EXTL1 Forward Primer: 5′-TGGGCACAGGAAGGTTAGTG-3′, EXTL1 Reverse Primer: 5′-CTTGTGGAAAGACTGCTGCG-3′, EXTL2 Forward Primer: 5′-ACTCGAGT GACAAGTGAGCC-3′, EXTL2 Reverse Primer: 5′-TGTGGCAACACCTCATTGTG-3′, EXTL3 Forward Primer: 5′-CAAGAAGTCGTGTGCTGAGG-3′, EXTL3 Reverse Primer: 5′-GCCAACAATCAGGCCATGTG-3′, GAPDH Forward Primer: 5′-TTGCCAT CAATGACCCCTTCA-3′, GAPDH Reverse Primer: 5′-CGCCCCACTTGATTTTGGA-3′.

#### BrdU proliferation assay

5×10^3^ HeLa shCTRL and shEXT1 cells were seeded in 96-well plate and cultured overnight. BrdU was added to the culture medium according to the manufacturer’s instructions (BrdU Cell Proliferation Assay Kit, Cell Signaling) and cells further incubated for 24, 48, and 72 h. The absorbance value for each well was measured at 450 nm with a microplate reader TECAN Infinite^®^200 PRO.

#### Proliferation assay using the xCelligence system

100 μl of cell culture media was added into each well of E-plate 96, and it was connected to the system in order the background impedance to be measured. HeLa shCTRL and shEXT1 cells were resuspended in cell culture medium and adjusted to 5.000 cells/well. 100 μl of each cell suspension was added to the 100 μl medium containing wells on E-plate 96. Cell index was monitored every 3 min for a period up to 72 h. The xCelligence system was used according to the instructions of the supplier Roche Applied Science and ACEA-Biosciences(2009).

#### Subcutaneous xenograft studies

Jurkat cells expressing control luciferase (shLUC or LUC-GFP) and the corresponding shEXT1 or EXT1-LUC-GFP, respectively were used. Briefly, 2⨯10^6^ viable human cells/type were mixed with an equal volume BD Matrigel^™^ basement membrane matrix and injected into the flanks of 6-week-old sublethally irradiated female NOD-SCID mice bred in-house and maintained under specific pathogen-free conditions. Cell growth and engraftment were monitored every 3 days (Caliper, Perkin Elmer). Animals were given an intraperitoneal injection of 150 mg/kg D-luciferin (Promega) and were imaged in groups of up to 3 mice (for display purposes).

#### Transmission electron microscopy

HeLa, HEK293 and Jurkat shCTRL and shEXT1, and HeLa shEXT2, shEXTL1, shEXTL2, shEXTL3 cells but also activated naive CD4^+^ T-cells from peripheral lymph organs (spleen and lymph nodes) of EXT1 F/F;LCK-CRE and LCK-CRE mice were fixed for 90 min at 4°C with 2.5% glutaraldehyde in Sörensen 0.1 M phosphate buffer (pH 7.4), and post-fixed for 30 min with 2% osmium tetroxide. Following, dehydration in graded ethanol, samples were embedded in Epon. Ultrathin sections obtained with a Reichert Ultracut S ultramicrotome were contrasted with uranyl acetate and lead citrate. The analysis was performed with a JEOL JEM-1400 transmission electron microscope at 80 kV and in a Tecnai Spirit T12 at 120 kV (Thermo Fisher Scientific, The Netherlands).

#### Immunohistochemistry

Immunohistochemical experiments were performed using a standard protocol previously described (Hubert et al., 2014). In the present study, the antigen retrieval step was: citrate pH 6.0 and the following primary antibody was used: anti-EXT1 (1/50, ab126305, Abcam,). The rabbit Envision kit (Dako) was used for the secondary reaction.

#### Flow cytometry, extracellular and intracellular staining

Single-cell suspensions from spleen, lymph node, blood, and thymus were prepared as described above. 1⨯10^6^ cells used for staining. Cells were resuspended in PBS and stained with the following fluorochrome-conjugated monoclonal antibodies purchased from (BD Biosciences): anti-mouse CD4-PE CF594 (RM4-5), anti-mouse CD8a-APC-H7 (53-6.7), anti-mouse CD45R/B220-V500 (RA3-6B2), anti-mouse CD25-BB515 (PC61) for 30 min at 4°C. Cells were washed twice and analyzed by FACS. Extracellular stains were performed in PBS supplemented with 0.5% BSA and 10% 24G.2 blocking antibody. After polarization, 1⨯10^6^ cells were fixed and permeabilized using Foxp3/Transcription Factor intracellular staining buffer set (eBioscience). The following conjugated mAbs were used: anti-mouse IL-17A-efluor450 (eBio17B7), anti-mouse IFN-γ-APC (XMG1.2) and anti-mouse IL-4-APC (11B11). The CFSE dye was used to label the dead cells. Cells analyzed immediately by flow cytometry on a BD LSRFortessa flow cytometer (BD Biosciences).

#### Plasmids

HA-SEC13 pRK5 (#46332), mEmerald-Sec61b-C1 (#90992) (Nixon-Abell et al., 2016), pEGFP-SEC16b (#66607) (Budnik et al., 2011), pEGFP-SEC23A (#66609) (Stephens et al., 2000), Str-KDEL-TNF-SBP-mCherry (#65279) (Boncompain et al., 2012), b4GALT1-pmTirquoise2-N1 (Goedhart et al., 2012) (#36205) constructs were obtained from Addgene. ts045-VSVG-GFP (#11912) (Presley et al., 1997) is a gift from Dr. Florian Heyd (Freie Universität Berlin, Berlin, Germany). EXT1-YFP and Flag-EXT1 were previously described (Daakour et al., 2016). Additional cloning vectors used here are: pCSCherryDEST (addgene#13075), mEmerald-C1 (addgene #53975) and pSYFP2-C1 (addgene #22878) (Kremers et al., 2006) or pCS2 EIF ires GFP. The lentiviral constructs used are: shCTRL (anti-eGFP, SHC005, Sigma-Aldrich) or pLV U6 shRNA NT PGK GFP-T2A-Neo and targeting EXT1 (sh438: TRCN0000039993, sh442: TRCN0000039997, Sigma-Aldrich). The shRNAs targeting EXT2, EXTL1, EXTL2, and EXTL3 were designed using Vector Builder online platform (https://en.vectorbuilder.com/) and cloned into lentiviral vector pLV-PURO-U6. The target sequences are listed below: EXT2: 5′-AGCGTACTTCCAGTCAATTAAC-3′ or 5′-CCATTGATGATATCATTA-3′, EXTL1: 5′-TGATCGCTTCTACCCATATAG-3′ or 5′-ATACCACTCTGGAGGTTATTC-3′; EXTL2: 5′-CTCTACTTCATCAGGTATCTA-3′ or 5′-GATTCGAGTGCTTCGATTATC-3′; EXTL3: 5′-CCGTACTGAGAAGAACAGTTT-3′ or 5′-TTGCCATTCAAGGCTTATTTA-3′. mCherry-RTN4a, mChery-ATL1, Lnp1-mCherry lentiviral constructs were a gift from Dr. Tom Rapoport (Dept of Cell Biology, Harvard Medical School, MA, USA). LV-PA-KDEL-GFP is a gift from Dr. Vicky C Jones (University of Central Lancashire, Preston, UK), Lenti-ATL3-GFP is a gift from Dr. Vincent Timmerman (University of Antwerp, Antwerp, Belgium). Lentivirus production and instructions on its use were kindly provided by Viral Vectors core facility (Viral Vectors platform, University of Liege).

#### Mammalian cell lines generation and culture

All cell lines HeLa, HEK293, Jurkat, and Cos7 were cultured as previously described (Daakour et al., 2016; Hu et al., 2009). All stable cell lines were generated by lentiviral transduction. Briefly, HEK293T Lenti-x 1B4 cells (Clontech®-Lenti-x HEK293T cells) were transfected with calcium phosphate with three plasmids: the vector of interest, pVSV-G (PT3343-5, Clontech) and psPAX2 (#12260, Addgene). The supernatants containing the second-generation viral vectors were harvested and concentrated by ultracentrifugation. The cells (HeLa, HEK293, Jurkat, Cos7) were transduced with the viral vector of interest with MOI (50, 80, 100 depending on the production). After 72 h, the cells were selected for puromycin (Invivogen) for 3-4 days. For fluorescence-protein-tagged constructs, positive cells were selected by flow cytometry sorting. The cells were finally tested for the presence of mycoplasma (MycoAlert Detection Kit, Lonza® LT07-318), and recombinant viral particles (Lentiviral qPCR TitrationKit, abmGood® #LV900).

#### DNA-siRNA transfection

DNA was transfected into HeLa and Cos7 with polyethylenimine (PEI 25K, Polysciences) as previously described (Daakour et al., 2016). For siRNA transfection, Cos7 and HeLa cells were transfected at 40-50% confluence with 2 nmol of siRNA using a classical calcium-phosphate method according to manufacturer’s instructions (ProFection Mammalian Transfection kit, Promega). The medium was changed 24 h later and cells were collected 48 h post-transfection. When experiments involved both DNA and siRNA transfection, siRNA transfection was performed, and 24 h later cells were transfected with DNA as described previously (Daakour et al., 2016). Cells were collected 24 h later. The following siRNA duplexes were purchased from Eurogentec (Belgium): siEXT1(1): 5′-GGAUCAUCCCAGGACAGGA -3′, siEXT1(2): 5′-GGAUUCCAGCGUGCACAUU-3′ and siCTRL: 5′-GGCUGCUUCUAUGAUUAUGtt-3′.

#### Calcium Flux Detection assay

2⨯10^5^ Cos7 cells were washed twice and processed for immunofluorescence. Fluo-4, AM Loading Solution was added on the cells according to manufacturer’s instructions (Fluo-4 Calcium Imaging Kit, Thermo Fisher Scientific). Images were acquired using a Leica TCS SP5 confocal microscope and the 63x oil objective; the analysis was performed in ImageJ software (Schindelin et al., 2012).

#### Preparation of microsomes from cultured cells

HeLa cells expressing FLAG-EXT1 or HeLa shCTRL and shEXT1 (2⨯10^8^) were harvested and washed with PBS and with a hypotonic extraction buffer (10 mM HEPES, pH 7.8, with 1 mM EGTA and 25 mM potassium chloride) supplemented with a protease inhibitors cocktail. Cells were resuspended in an isotonic extraction buffer (10 mM HEPES, pH 7.8, with 0.25 M sucrose, 1 mM EGTA, and 25 mM potassium chloride) supplemented with a protease inhibitors cocktail and homogenized with 10 strokes using a Dounce homogenizer. The suspension was centrifuged at 1.000 g for 10 min at 4°C. The supernatant was centrifuged at 12.000 g for 15 min at 4°C. The following supernatant fraction, which is the post mitochondrial fraction (PMF), is the source for microsomes. The PMF was centrifuged for 60 min at 100.000 g at 4°C. The pellet was resuspended in isotonic extraction buffer supplemented with a protease inhibitors cocktail and stored in −80°C. Isolated membranes were boiled 5 min in 2x SDS-loading buffer. Then, solubilized samples were separated on SDS-PAGE and analyzed by western blotting.

#### Western blotting and antibodies

Cells were lysed in immunoprecipitation low salt buffer (IPLS: 25 mM Tris-HCl pH 7.4, 150 mM NaCl, 1 mM EDTA, 1% NP-40 and 5% glycerol, complete Protease Inhibitor (Roche) and Halt Phosphatase Inhibitors (Thermo Fisher Scientific)). Concentrations were determined using the Bradford assay. SDS-PAGE and western blotting were performed using standard protocols. The following primary antibodies were used: mouse-anti-Calnexin 1:2000 (Abcam), rabbit-anti-EXT1 1:500 (Prestige Antibodies, Sigma-Aldrich), mouse-anti-NogoA (Santa Cruz), rabbit-anti-FLAG 1:4000 (Sigma-Aldrich), mouse-anti-FLAG 1:4000 (Sigma-Aldrich), goat-anti-actin 1:2000 (Santa Cruz), rabbit-anti-HSP70 1:3000 (Santa Cruz). Dad1, STT3b, STT3a, Sec61A, Trap-alpha, TRAP-beta, SEC62, SEC63 rabbit antibodies were a kind gift from Dr. Richard Zimmermann (Medical Biochemistry and Molecular Biology, Saarland University, Homburg, Germany). The following conjugated secondary antibodies were used: a-mouse-HRP 1:5000 (Santa Cruz), a-rabbit-HRP 1:5000 (Santa-Cruz), anti-goat 1:5000 (Santa-Cruz).

#### N-glycans and O-glycans profiling

Microsomes were isolated as described above, and glycans profiling performed by Creative Proteomics (NY, USA). For the preparation of *N*-glycans ∼250 µg of lyophilized protein samples are required. The dry samples are resuspended in fresh 2 mg/ml solution of 1,4-dithiothreitol in 0.6 M TRIS buffer pH 8.5 and incubated at 50°C for 1 h. Fresh 12 mg/ml solution of iodoacetamide in 0.6 M TRIS buffer pH 8.5 was added to the DTT-treated samples and incubated at RT in the dark for 1 h. Samples were dialyzed against 50 mM ammonium bicarbonate at 4°C for 16-24 h, changing the buffer 3 times. The molecular cut-off should be between 1 and 5 kDa. After dialysis, the samples were transferred into 15 ml tubes and lyophilized. Following resuspension of the dry samples in 0.5 ml of a 50 µg/ml solution of TPCK-treated trypsin in 50 mM ammonium bicarbonate and overnight incubation at 37°C. The reactions stopped by adding 2 drops of 5% acetic acid. Condition a C18 Spe-Pak (50 mg) column with methanol, 5% acetic acid, 1-propanol and 5% acetic acid. Trypsin-digested samples were loaded onto the C18 column and then column was washed with 4 ml of 5% acetic acid and the peptides eluted from the C18 column with 2 ml of 20% 1-propanol, then 2 ml 40% 1-propanol, and finally 2 ml of 100% isopropanol. All the eluted fractions were pooled and lyophilized. The dried material was resuspended thoughtfully in 200 µl of 50 mM ammonium bicarbonate and 2 µl of PNGaseF was added, following incubation at 37°C for 4 h. Then, another 3 µl of PNGaseF was added for overnight incubation at 37°C. To stop the reaction addition of 2 drops of 5% acetic acid is required. Condition a C18 Spe-Pak (50 mg) column with methanol, 5% acetic acid, isopropanol and 5% acetic acid and the PNGaseF-digested samples were loaded onto the C18 column, and flow-through was collected. The column was washed with 4 ml of 5% acetic acid, and fractions were collected. Flow-through and wash fractions were pooled, samples were lyophilized and proceeded to permethylation.

For the *O*-glycans preparation, 1 ml of 0.1 M NaOH was added to 55 mg of NaBH4 in a clean glass tube and mixed well, and 400 μl of the borohydride solution was added to the lyophilized sample (collected peptides/glycopeptides after PNGaseF digestion). Following, incubation at 45°C overnight, the reaction was terminated by the addition of 4-6 drops of pure (100%) acetic acid, until fizzing stops. A stock solution of Dowex 50W X8 (mesh size 200-400) was made by washing three times 100 g of resin with 100 ml of 4 M HCl. The resin was washed with 300 ml of Milli-Q water, and the wash step was repeated for ∼15 times until the pH remained stable. The resin was then washed with 200 ml of 5% acetic acid three times. A desalting column with 2-3 ml of the Dowex resin prepared above in a small glass column. The column was washed with 10 ml of 5% acetic acid. Acetic acid-neutralized samples were loaded onto the column and washed with 3 ml of 5% acetic acid. Flow-through was pooled and washed. The collected material was lyophilized, supplemented with 1 ml of acetic acid: methanol (1:9;v/v=10%) solution, vortexed thoroughly and dried under a stream of nitrogen. This co-evaporation step was repeated for three more times. Condition a C18 Spe-Pak column with methanol, 5% acetic acid, isopropanol and 5% acetic acid. The dried sample was resuspended in 200 μl of 50% methanol and loaded onto the conditioned C18 column. The column was washed with 4 ml of 5% acetic acid. Flow-through was collected, pooled, and washed. Lyophilized samples were processed to permethylation.

For the permethylation, the preparation of the slurry NaOH/DMSO solution is made fresh every time. Mortar, pestle, and glass tubes were washed with Milli-Q water and dried beforehand. Whenever possible, liquid reagents were handled with disposable glass pipettes. Solvents are HPLC grade or higher. With a clean and dry mortar and pestle grind 7 pellets of NaOH in 3 ml of DMSO.

One ml of this slurry solution was added to a dry sample in a glass tube with a screw cap and supplemented with 500 µl of Iodomethane and incubated at RT for 30 min. The mixture turns white and even becomes solid as it reaches completion. One ml of Milli-Q water was added to stop the reaction, and the tube was vortexed until all solids were dissolved. The sample was supplemented with 1 ml of Chloroform and additional 3 ml of Milli-Q water, vortexed and centrifuged briefly to separate the chloroform and the water phases (∼5.000 rpm, <20 sec). The aqueous top layer was discarded and wash 2 more times. Chloroform fraction dried with a SpeedVac (∼20-30 min). Condition a C18 Spe-Pak (200 mg) column with methanol, Milli-Q water, and acetonitrile. Dry samples were resuspended in 200 µl of 50% methanol and loaded onto the column. The tube was washed with 1 ml of 15% acetonitrile and loaded onto the column. The column was washed with 2 ml of 15% acetonitrile, then eluted in a clean glass tube with 3 ml of 50% acetonitrile. Lyophilized eluted fraction for MS analysis was used. MS data were acquired on a Bruker UltraFlex II MALDI-TOF Mass Spectrometer instrument. The positive reflective mode was used, and data were recorded between 500 m/z and 6000 m/z for *N*-glycans and between 0 m/z and 5000 m/z for *O*-glycans. For each MS *N*- and *O*-glycan profiles the aggregation of 20.000 laser shots or more were considered for data extraction. Mass signals of a signal/noise ratio of at least 2 were considered and only MS signals matching an *N*- and *O*-glycan composition was considered for further analysis and annotated. Subsequent MS post-data acquisition analysis was made using mMass(Strohalm et al., 2010).

#### Glycosyltransferase assay

Glycosyltransferase activity of microsomes from HeLa shCTRL, and shEXT1 was determined with the Glycosyltransferase Activity Kit (R&D Systems). A glycosyltransferase reaction was carried out in 50 µL of reaction buffer in a 96-well plate at room temperature for 20 min, according to the manufacturer’s instructions. The absorbance value for each well was measured at 620 nm with a microplate reader TECAN Infinite®200 PRO.

#### Metabolomics profiling

For metabolite quantification, HEK293 shCTRL, and shEXT1 cells were seeded in triplicate (n=3) in 6-well plates with DMEM supplemented with 10% FBS. After 24 h, the media was removed and replaced with fresh media containing stable isotopic tracer ^13^C-glucose. For one well per condition, the medium was replaced with 1-^12^C-glucose. Upon reaching 70% confluency, the supernatant was stored in −80°C and cells were washed twice with PBS, harvested and the cell pellet stored in −80°C until Liquid Chromatography/Mass Spectrometry identification of metabolites at the University of Leuven metabolomics core facility.

#### Lipidomics

20 μg protein or ER microsomes diluted in 700 μl water was mixed with 800 μl 1 N HCl:CH3OH 1:8 (v/v) and 900 μl CHCl3, in the presence of 200 μg/ml of the antioxidant 2,6-di-tert-butyl-4-methylphenol (BHT; Sigma Aldrich). 3 μl of SPLASH® LIPIDOMIX® Mass Spec Standard (#330707, Avanti Polar Lipids) was spiked into this mixture and the samples were vortexed and centrifuged at 4000g for 10 min. The organic fraction was evaporated using a Savant Speedvac spd111v (Thermo Fisher Scientific) at room temperature and the remaining lipid pellet was stored at −20°C under argon. Just before mass spectrometry analysis, lipid pellets were reconstituted in 100% ethanol. Lipid species were analyzed by hydrophilic interaction liquid chromatography electrospray ionization tandem mass spectrometry (HILIC-ESI/MS/MS) on a Nexera X2 UHPLC system (Shimadzu) coupled with a hybrid triple quadrupole/linear ion trap mass spectrometer (6500+ QTRAP system; AB SCIEX). Chromatographic separation was performed on a XBridge amide column (150 mm × 4.6 mm, 3.5 μm; Waters) maintained at 35°C using mobile phase A [1 mM ammonium acetate in water-acetonitrile 5:95 (v/v)] and mobile phase B [1 mM ammonium acetate in water-acetonitrile 50:50 (v/v)] using the gradient (0-6 min: 0% B → 6% B; 6-10 min: 6% B → 25% B; 10-11 min: 25% B → 98% B; 11-13 min: 98% B → 100% B; 13-19 min: 100% B; 19-24 min: 0% B) at a flow rate of 0.7 mL/min which was increased to 1.5 mL/min from 13 minutes onwards. Lipid quantification was performed by scheduled multiple reactions monitoring (MRM), the transitions being based on the generation of neutral losses or typical fragment ions during collision induced dissociation in tandem MS. SM, CE, CER, DCER, HCER, LCER were measured in positive ion mode with MRM transitions based on the generation of fragment ions of m/z 184.1, 369.4, 264.4, 266.4, 264.4 and 266.4, respectively. TAG, DAG and MAG were measured in positive ion mode with MRM transitions based on the neutral loss of each of the fatty acyl moieties. PC, LPC, PE, LPE, PG, LPG, PI, LPI, PS and LPS were measured in negative ion mode with MRM transitions based on the neutral loss of each of the fatty acyl moieties. The instrument parameters were as follows: Curtain Gas = 35 psi; Collision Gas = 8 a.u. (medium); IonSpray Voltage = 5500 V/ −4,500 V; Temperature = 550°C; Ion Source Gas 1 = 50 psi; Ion Source Gas 2 = 60 psi; Declustering Potential = 60 V/ −80 V; Entrance Potential = 10 V/ −10 V; Collision Cell Exit Potential = 15 V/ −15 V. Lipidomics analysis was performed in the Lipidomics core facility of the University of Leuven, Laboratory of Lipid Metabolism.

#### Immunofluorescence and confocal, super-resolution microscopy

3⨯10^4^ Cos7 and 5⨯10^4^ HeLa cells were grown on 18 mm round glass coverslips and transfected with 500 ng of DNA/well. For immunostaining, the cells were washed with PBS (pH 7.4) and fixed with 4% paraformaldehyde in PBS for 15 min at RT. Cells were permeabilized with 0.5% Triton X-100 for 10 min and incubated with blocking solution (0.025% Tween-20 and 10% FBS) for 30 min. Primary antibody staining was performed overnight at 4°C in 5% blocking solution: mouse-anti-beta-catenin 1:1000 (Santa Cruz, RRID:AB_626807), mouse-anti-Calnexin 1:500 (Abcam, RRID:AB_2069009), rabbit-anti-EXT1 1:100 (Prestige Antibodies Sigma-Aldrich, RRID:AB_10963838), mouse-anti-HS (10E4) (1:100, USBio, RRID:AB_10013601), rabbit-anti-GM130 1:3200 (Cell Signaling, RRID:AB_2797933), mouse-anti-PDIA3 1:1000 (Prestige Antibodies Sigma-Aldrich, RRID:AB_2665750), mouse-anti-SEC31 1:500 (BD Bioscience, RRID:AB_399716). Goat-anti-rabbit, donkey-anti-rabbit or goat-anti-mouse secondary antibodies labeled with Alexa Fluor 488 or Texas Red (Thermo Fisher Scientific), anti-mouse-STAR-Red (Abberior) were used at a 1:2000 dilution for 1 h. Cells were stained with DAPI (Thermo Fisher Scientific) when needed for 5 min at RT, washed 5 times with PBS and mounted with Prolong Antifade Mountants (Thermo Fisher Scientific). Slides were analyzed by confocal microscopy with a Leica TCS SP8 microscope using the 100x oil objective. Images were taken at 2068 × 2068 pixel resolution and deconvoluted with Huygens Professional software. SYFP2-EXT1 was analyzed by Stimulated Emission Depletion (STED) microscopy with a Leica SP8 STED 592 nm laser. Images were taken at 2068 × 2068 pixel resolution and deconvoluted with Huygens Professional software. SEC31 was analyzed with Stedycon STED laser 775 nm. mEmerald-EXT1 was analyzed by Structured Illumination Microscopy (SIM) super-resolution. SIM imaging was performed at the Cell Imaging and Cytometry Core facility (Turku University) using a DeltaVision OMX SR V4 microscope using a 60x/1.42 Olympus Plan Apo N SIM objective and sCMOS cameras (Applied Precision), 2560 ⨯ 2160 pixel resolution. The SIM image reconstruction was performed with DeltaVision softWoRf 7.0 software. For live imaging of Cos7 cells expressing mCherry-ATL1 or Lnp1-mCherry, 3⨯10^4^ cells were plated and imaged at 37°C and 5% CO_2_ in a thermostat-controlled chamber on a Zeiss LSM800 AiryScan Elyra S1 SR confocal microscope using the 63x oil objective at 1 frame/100 ms for 5 s. Further analysis was performed in ImageJ software (Schindelin et al., 2012).

#### Photoactivatable GFP Imaging

Using an adaptation of a published assay (Krols et al., 2018), 3⨯10^4^ Cos7 cells expressing PA-KDEL-GFP were plated, and live imaging was performed at 37°C and 5% CO_2_ in a thermostat-controlled chamber on a Zeiss LSM800 AiryScan Elyra S1 SR confocal microscope using the 100x oil-objective. PA-KDEL-GFP was activated at a perinuclear ER region using the 405 nm laser at 100%, after which the cell was imaged at 1 frame/500 ms for 90 s using the 488 nm laser. Fluorescence intensities were measured using ImageJ software (Schindelin et al., 2012), and data analysis and curve fitting were performed in Graphpad Prism 8 (Graphpad Software). To avoid inter-cell variability, the activation site was at the perinuclear area of cells with the same ER density. The integrated fluorescence intensity of each region of interest (ROI) at fixed distances (8,12,16 μm) from the activation region was measured in ImageJ. Normalization of raw values was done, by defining the initial fluorescence to zero and the maximum fluorescence to 1 for each ROI. Image analysis was performed in ImageJ (Schindelin et al., 2012).

#### Affinity purification for mass spectrometry

2x solubilization buffer (3.5% digitonin, 100 mM HEPES (pH 7.5), 800 mM KOAc, 20 mM MgOAc2, 2 mM DTT) was mixed in a ratio 1:1 with the microsomal fraction and incubated 10 min on ice. Samples were centrifuged for 15 min at 14.000 rpm to isolate the solubilized material and remove the insoluble material. The supernatant was further used for immunoprecipitation. Equilibrated agarose beads M2-FLAG (Sigma-Aldrich) were added in the microsomal fraction (15 µl of beads per half of a 10-cm cell culture dish), and rotation was performed overnight at 4°C. Beads were washed 3 times for 15 min with glycine 50 mM pH 3.0 for protein elution. The supernatant was supplemented with Tris-HCL pH 8.0. Eluted proteins were then subjected to trypsin digestion and identified by mass spectrometry. Mass spectrometry analyses were performed by the GIGA-Proteomics facility, University of Liege or the proteomic core facility of de Duve Institute, Brussels, Université Catholique de Louvain, Belgium.

As a control, beads were washed five times with IPLS and eluted by boiling 5 min in 2x SDS-loading buffer. Then, solubilized samples were separated on SDS-PAGE and analyzed by western blotting.

#### Mass spectrometry

Peptides were dissolved in solvent A (0.1% TFA in 2% ACN), directly loaded onto reversed-phase pre-column (Acclaim PepMap 100, Thermo Fisher Scientific). Peptide separation was performed over 140 min using a reversed-phase analytical column (Acclaim PepMap RSLC, 0.075 ⨯ 250 mm, Thermo Fisher Scientific) with a linear gradient of 4%-32% solvent B (0.1% FA in 98% ACN) for 100 min, 32%-60% solvent B for 10 min, 60%-95% solvent B for 1 min and holding at 95% for the last 6 min at a constant flow rate of 300 nl/min on an Ultimate 3000 UPLC system. The resulting peptides were analyzed by Orbitrap Fusion Lumos tribrid mass spectrometer using a high-low data-dependent scan routine for protein identification and an acquisition strategy termed HCD product-dependent EThcD/CID (Thermo Fisher Scientific) for glycopeptides analysis.

Briefly for the latter, the peptides were subjected to NSI source and were detected in the Orbitrap at a resolution of 120.000. Peptides were selected for MS/MS using HCD setting as 28 and detected in the Orbitrap at a resolution of 30.000. If predefined glycan oxonium ions were detected in the low m/z region it triggered an automated EThcD and CID spectra on the glycopeptide precursors in the Orbitrap. A data-dependent procedure that alternated between one MS scan every 3 seconds and MS/MS scans was applied for the top precursor ions above a threshold ion count of 2.5E4 in the MS survey scan with 30.0s dynamic exclusion. MS1 spectra were obtained with an AGC target of 4E5 ions and a maximum injection time of 50 ms, and MS2 spectra were acquired in the Orbitrap at a resolution of 30.000 with an AGC target of 5E4 ions and a maximum injection time of 300 ms. For MS scans, the m/z scan range was 350 to 1800. For glycopeptide identification the resulting MS/MS data was processed using Byonic 3.5 (Protein Metrics) search engine within Proteome Discoverer 2.3 against a human database obtained from Uniprot, the glycan database was set to “*N*-glycan 182 human no multiple fucose or *O*-glycan 70 human”. Trypsin was specified as cleavage enzyme allowing up to 2 missed cleavages, 5 modifications per peptide and up to 7 charges. Mass error was set to 10 ppm for precursor ions and 20 ppm for fragment ions. Oxidation on Met, carbamidomethyl (+57.021 Da) were considered as variable modifications on Cys. Glycopeptides with a Byoinic score >= 300 and with a Log Prob >= 4.0 were retained and their identification was manually validated.

#### SILAC labeling

HeLa cells (shCTRL, shEXT1) were cultured for at least five cell doublings in either isotopically light or heavy SILAC DMEM obtained from Thermo Scientific (catalog number A33969) containing 10% FBS and 50 μg/ml streptomycin and 50 units/ml penicillin (Lonza). For the heavy SILAC medium, 50 mg of ^13^C_6_ L-Lysine-2HCl (heavy) and 50 mg of L-Arginine-HCl was added. In light SILAC medium 50 mg of L-Lysine-2HCl (light) and 50 mg of L-Arginine-HCl was added. 2×10^5^ cells adapted to grow in DMEM. The cell pellet was suspended in 150 μl of modified RIPA buffer and sonicated followed by incubation at 60°C for 15 min. Samples were clarified by centrifugation; each replicate was pooled and quantified by Qubit (Invitrogen): 20 μg of the sample was separated on a 4-12% Bis-Tris Novex mini-gel (Invitrogen) using the MOPS buffer system. The gel was stained with coomassie, and gel bands were excised at 50 kDa and 100 kDa. Gel pieces were processed using a robot (ProGest, DigiLab). They were washed with 25 mM ammonium bicarbonate followed by acetonitrile and reduced with 10 mM dithiothreitol at 60°C followed by alkylation with 50 mM iodoacetamide at RT and digested with trypsin at 37°C for 4 h. Finally, they were quenched with formic acid, and the supernatant was analyzed directly without further processing. For the SILAC analysis performed by MS Bioworks LLC (MI, USA), the samples were pooled 1:1 and 20 μg was separated on a 4-12% Bis-Tris Novex minigel (Invitrogen) using the MOPS buffer system. The gel was stained with coomassie, and the lanes excised into 40 equal segments using a grid. For mass spectrometry, the gel digests were analyzed by nano-LC/MS/MS with a Waters NanoAcquity HPLC system interfaced to a Thermo Fisher Q Exactive. Peptides were loaded on a trapping column and eluted over a 75 μm analytical column at 350 nL/min. Both columns were packed with Luna C18 resin (Phenomenex). The mass spectrometer was operated in data-dependent mode, with MS and MS/MS performed in the Orbitrap at 70.000 FWHM and 17.500 FWHM resolution, respectively. The fifteen most abundant ions were selected for MS/MS. Data were processed through the MaxQuant software 1.5.3.0 (www.maxquant.org) which served several functions such as the recalibration of MS data, the filtering of database search results at the 1% protein and peptide false discovery rate (FDR), the calculation of SILAC heavy: light ratios and data normalization. Data were searched using a local copy of Andromeda with the following parameters, Enzyme set as trypsin, database set as Swissprot Human (concatenated forward and reverse plus common contaminant proteins), fixed modification: Carbamidomethyl (C), variable modifications: Oxidation (M), Acetyl (Protein N-term), 13C6 (K) and fragment Mass Tolerance: 20 ppm.

#### Rush assay

An adaptation of published assay (Boncompain et al., 2012) was used. HeLa cells were transfected with Str-KDEL-TNF-SBP-mCherry construct as described above, and 24 h after transfection mCherry positive cells were sorted. 5×10^4^ cells were cultured on 35 mm imaging dish. The day after, cells were transferred at 37°C in a thermostat-controlled chamber. At time point zero, the medium was removed and replaced with medium containing D-biotin (Sigma-Aldrich) at 40 μM concentration. The time-lapse acquisition was made using a Zeiss LSM800 AiryScan Elyra S1 SR confocal microscope. Images were acquired using a 63x oil-objective. For each time point, the integrated intensity of a region of interest (ROI) was measured. The integrated intensity of an identical size ROI corresponding to background was measured and subtracted from the values of the integrated intensity for each time point. The values were then normalized to the maximum value. These quantifications were performed using the Zeiss Black software.

#### Export assay

3⨯10^4^ Cos7 cells were cultured on 35 mm imaging dish, and transfected with the ts045-VSVG-GFP reporter construct and immediately incubated at 40°C overnight to retain the reporter protein in the ER. After the addition of cycloheximide, cells were transferred in a thermostat-controlled chamber at 40°C. The temperature was shifted to 32°C, and cells were processed for immunofluorescence at t=0, t=45 and t=90 min and stained with mouse-anti-beta-catenin antibody as described above. The acquisition was made using a Zeiss LSM800 AiryScan Elyra S1 SR confocal microscope. Images were acquired using a 40x oil-objective.

### QUANTIFICATION AND STATISTICAL ANALYSES

#### Image analysis

For colocalization analysis, the average Pearson’s correlation coefficient test was performed with the plugin Colocalization Threshold in ImageJ software (Schindelin et al., 2012).

To track the displacement of main junctions during successive frames, the dynamic features of the cell were retrieved from the time-lapses of Cos7 cells expressing mCherry-ATL1 or Lnp1-mCherry with the following image processing procedure. Images were pre-processed to uniformize the intensities. Then, each image was binarized and skeletonized using Matlab2016a. The skeleton was labeled using AnalyzeSkeleton plugin from ImageJ. From this process, each pixel of the skeleton was classified according to its neighborhood leading to three-pixel classes: end-point, junctions and tubules. To reflect the structure of the ER, the ratio of the junctions over the tubules was computed for mCherry-ATL1 and Lnp1-mCherry proteins. The dynamics of the ER was assessed by the main junctions displacement during a time-lapse. To achieve the tracking of the displacement, the junctions larger than three pixels were kept segmented. Then, the segmented objects were multiplied by the initial image intensity to consider the initial light intensity. Finally, a gaussian blur was applied to these objects. The tracking of the bright spot was achieved by using a single-particles tracking algorithm, the “simple LAP tracker” available in ImageJ plugin TrackMate (Tinevez et al., 2017). The parameters were set following the recommendations for Brownian motion like’s movements, i.e., a max linking distance of seven pixels, a max closing distance of ten pixels and a max frame gap of three pixels. From the results of Trackmate, only the tracks longer than ten frames were kept in order to reduce the noise. Finally, using all velocity vectors measured, a cumulative velocity distribution was computed. Furthermore, a diffusion coefficient based on instantaneous velocity was computed using the Matlab as described previously (Holcman et al., 2018).

In AnalyzER (Pain et al., 2019), original images were imported, and the regions of interest segmented using Otsu’s method (Otsu, 1979). Cisternae are identified using an image opening function and active contour refinement. The tubular network is enhanced using phase congruency, and the resulting enhanced network is skeletonized to produce a single-pixel wide skeleton running along each tubule. Regions fully enclosed by the skeletonized tubular network and the cisternae are defined as polygonal regions, and features such as area, circularity, and elongations are extracted.

#### SAFE analysis

We used the SAFE software (Baryshnikova, 2016) (v1.5) to determine and visualize significant functional modules i) in the network of EXT1 partners and their first-order neighbors excerpted from STRING (Szklarczyk et al., 2015) database with confidence over 0.95 ii) in the network of genes whose expressions are significantly regulated by EXT1 knockdown obtained from STRING database with confidence over 0.9. The network layouts were generated with Cytoscape (Shannon et al., 2003) (v3.4.0) using the edge-weighted spring embedded layout. Gene Ontology (Ashburner et al., 2000) (GO) terms for each gene were extracted from FuncAssociate (Berriz et al., 2003) (v3 - GO updated on February 2018). The SAFE analysis was run with the default option.

#### RNA sequencing

RNA sequencing analysis was previously described are deposited as GSE138030.

#### Model generation and flux balance analysis

Model generation and *in silico* flux balance analysis was done using the Constraint-Based Reconstruction Analysis (COBRA) toolbox V3.0 (Heirendt et al., 2019) in the Matlab 2018a environment with an interface to IBM Cplex and GLPK solvers provided in the COBRA toolbox. Linear programing problems were solved on a macOS Sierra version 10.12.6. To generate the control and EXT1 knocked down specific models, the gene expression mRNA data for samples of control EXT1 knocked down cells (RNA seq) were integrated with the COBRA human model, *RECON2* (Thiele et al., 2013). The integration step uses the GIMME algorithm (Becker and Palsson, 2008), available in the COBRA toolbox. Because GIMME requires binary entries for the indication of the presence or absence of genes, we used a gene expression threshold value equals to the first quartile RPKM (reads per kilobase of transcript per million) for genes in control and EXT1 knocked-down cells. GIMME only integrates reactions associated with active genes, leaving those associated with the lowly expressed genes inactive. Therefore, genes with expression values below the threshold were given the value of 0 (inactive), and those with expression values higher than the threshold were given a value of 1 (active). Flux balance analysis (FBA) calculates the flow of metabolites through a metabolic network, thereby predicting the flux of each reaction contributing to an optimized biological objective function such as growth rate. Simulating growth rate requires the inclusion of a reaction that represents the production of biomass, which corresponds to the rate at which metabolic precursors are converted into biomass components, such as lipids, nucleic acids, and proteins. For both models generated after the integration step, we used the biomass objective function as defined in the *RECON2* model to obtain the FBA solution using the COBRA Toolbox command, *optimizeCbModel*. After identification of the objective function in the model, the entries to the command *optimizeCbModel* are: the model and the required optimization of the objective function (maximum production). The command output is the FBA solution, which includes the value of the maximum production rate of the biomass and a column vector for the conversion rate value (reaction fluxes) of each metabolite accounted for in the model.

#### Statistical analysis

Graph values are represented as mean + s.d. (standard deviation) of the mean calculated on at least three independent experiments/samples. The analyses were performed in Prism 8 (Graphpad Software). The statistical significance between means was determined using one-way ANOVA followed by two-tailed, unpaired Student’s t-test. p-values thresholds depicted as follows: *p<0.05, **p<0.01, ***p<0.001, ****p<0.0001, n.s., not significant. Significance for PA-KDEL-GFP was performed using two-way ANOVA followed by Sidak’s multiple comparisons test. Significance for Rush assay was performed using the two-stage linear step-up procedure of Benjamini, Krieger and Yekutieli, with Q=1%. Each time point was analyzed individually, without assuming a consistent SD.

## DATA AVAILABILITY

The mass spectrometry proteomics data have been deposited to the ProteomeXchange Consortium via the PRIDE (Perez-Riverol et al., 2019) partner repository with the dataset identifier PXD015660 and 10.6019/PXD015660.

RNA-sequencing data have been deposited in NCBI’s Gene Expression Omnibus (Edgar, 2002) and are accessible through GEO accession number GSE138030 (https://www.ncbi.nlm.nih.gov/geo/query/acc.cgi?acc=GSE138030). Supporting datasets are available at 10.17632/2mfzds3mmv.1 and 10.17632/y3h34szx5z.1.

## SUPPLEMENTAL EXCEL TABLES

**Table S1. Related to Figures 2, 3 and 4**

Quantification of ER extension, Cell size, Golgi apparatus size, secretion vesicles and ER-mitochodria/nuclear enveloppe contact sites.

**Table S2. Related to Figure 5**

N- and O-glycans on ER membrane proteins from HeLa *EXT1 k*.*d*. and control cells.

**Table S3. Related to Figure 5**

Analysis of glycosylation of OST catalytic subunits in ER microsomes from HeLa *EXT1 k*.*d*. and control cells.

**Table S4. Related to Figure 5**

Proteomic analysis of ER membrane proteins from HeLa EXT1 k.d. and control cells.

**Table S5. Related to Figure 5**

Quantification of lipid classes in ER microsomes from HeLa *EXT1 k*.*d*. and control cells.

**Table S6. Related to Figure 7**

Proteins identified by mass spectrometry, as partners of EXT1 in ER membranes.

**Table S7. Related to Figure 7**

SILAC analysis of HeLa *EXT1 k*.*d*. and control cells.

