## Supplemental Figure 1 for "Exostosin-1 Glycosyltransferase Regulates Endoplasmic Reticulum Architecture and Dynamics"

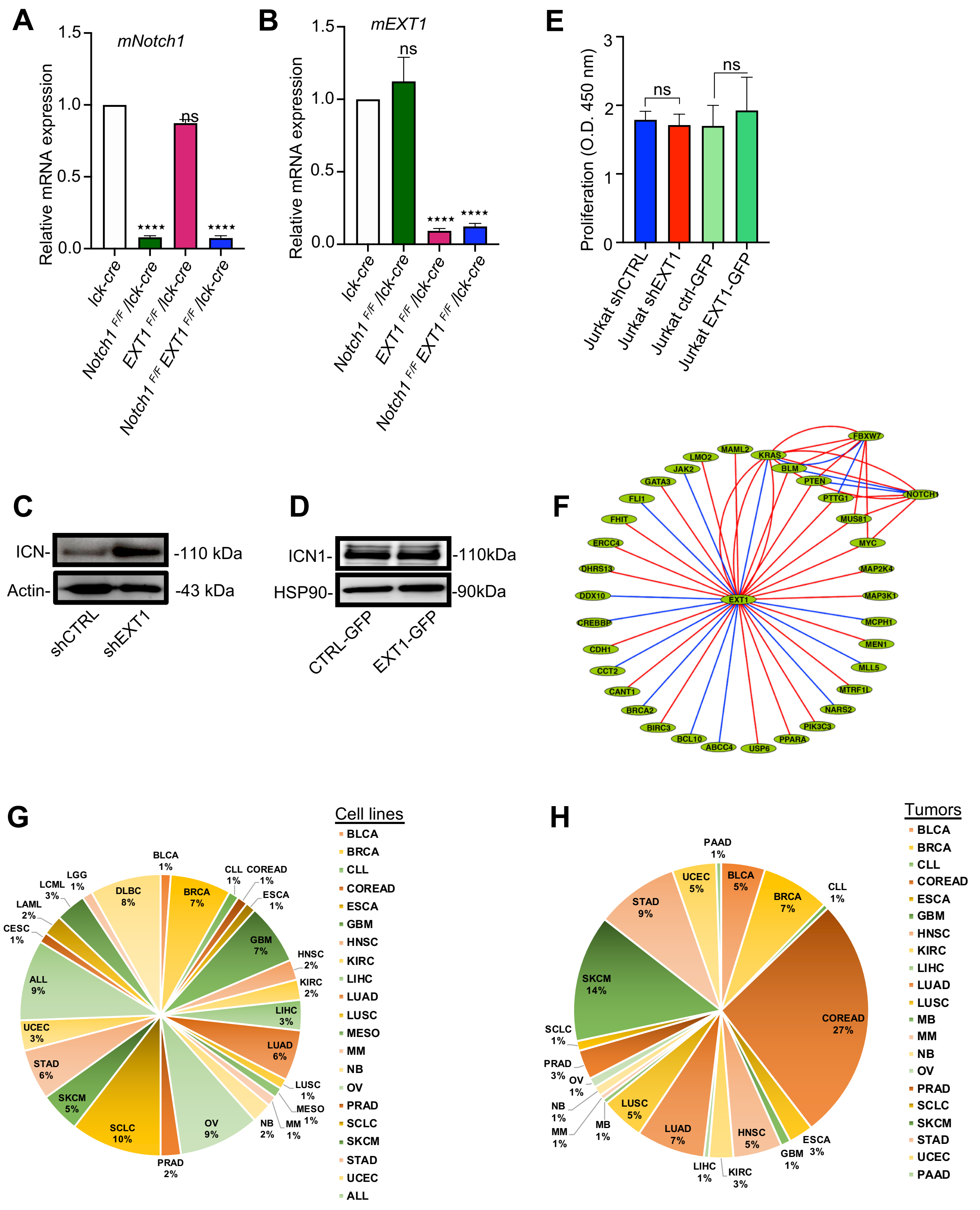


**Figure S1. *EXT1* is a suppressor hub in different cancer types, Related to Figures 1 and 2**

**(A-B)** Relative mRNA expression levels of mouse *EXT1* (A) and mouse *Notch1* (B) in thymocytes isolated from indicated mice. One-way ANOVA: ****p<0.0001; ns: not significant. **(C-D)** Western blot analysis of intracellular Notch1 (ICN1) in Jurkat-luciferase shEXT1 (C) or Jurkat-luciferase EXT1-GFP (D) compared to control cells. **(E)** Proliferation of Jurkat cells measured with BrdU optical absorbance at 450 nm. Mean number + SD is plotted. One-way ANOVA: ns: not significant. **(F)** Network depicting the synthetic lethality (SL) interactome map of EXT1. Blue edges indicate SL observed experimentally and verified clinically, while red edges indicate SL not verified clinically. **(G-H)** Pie chart representing the percentage of EXT1 mutations in cancer cell lines (G) and in tumor samples (H). Data are from TCGA and COSMIC databases.
