## Supplementary figures and images for "Exostosin-1 Glycosyltransferase Regulates Endoplasmic Reticulum Architecture and Dynamics"

### Supplemental Figure 2

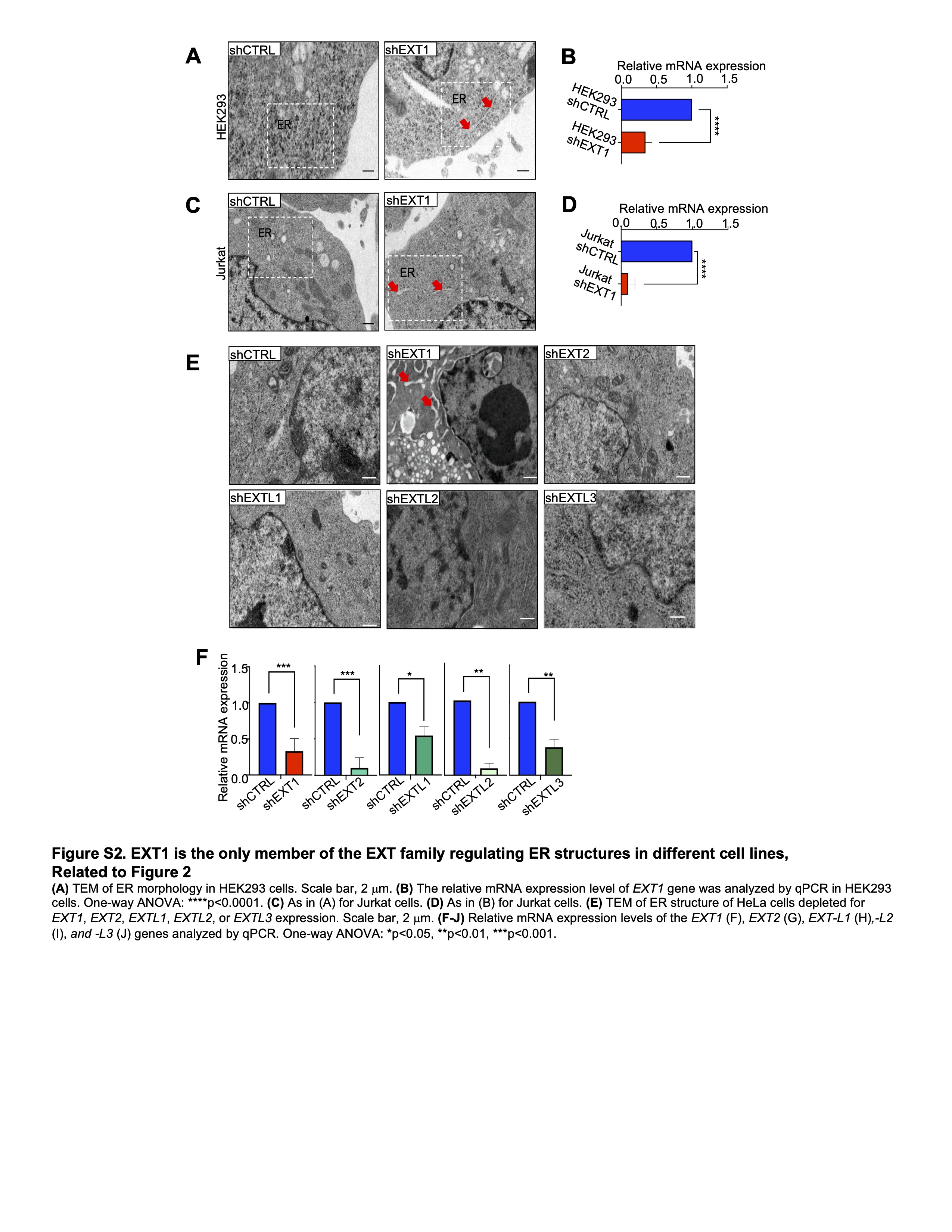

### Supplemental Figure 5

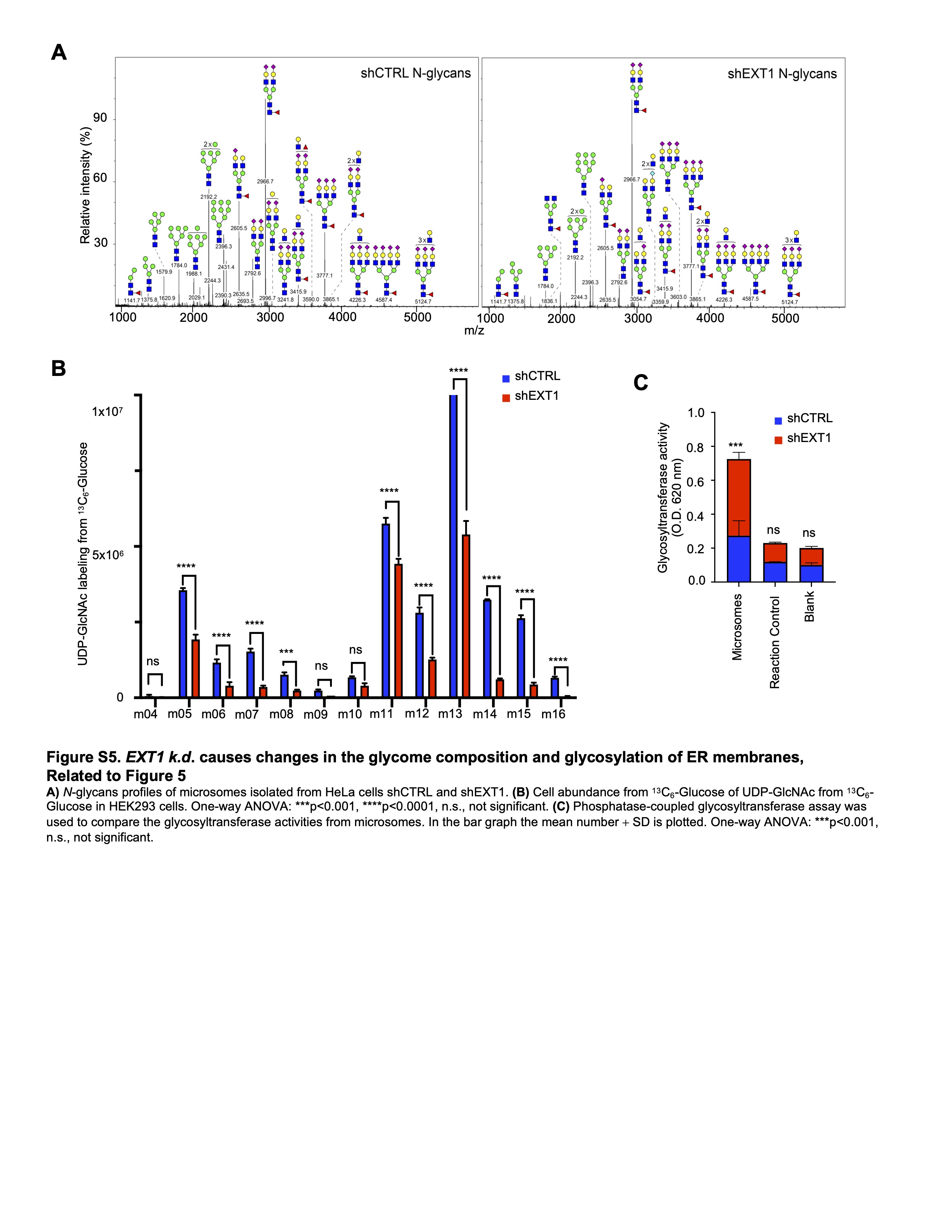

### Supplemental Figure 6

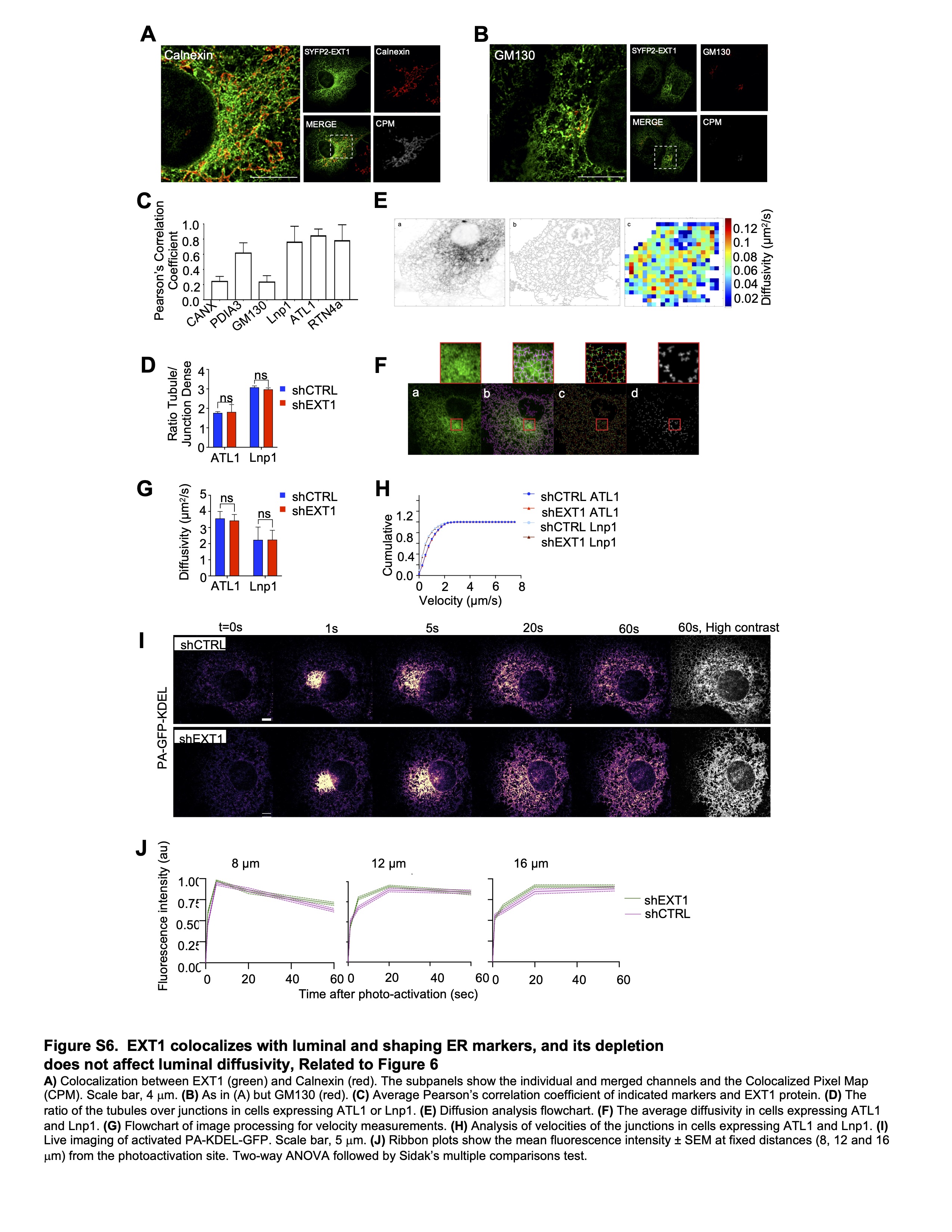

### Supplemental Figure 7

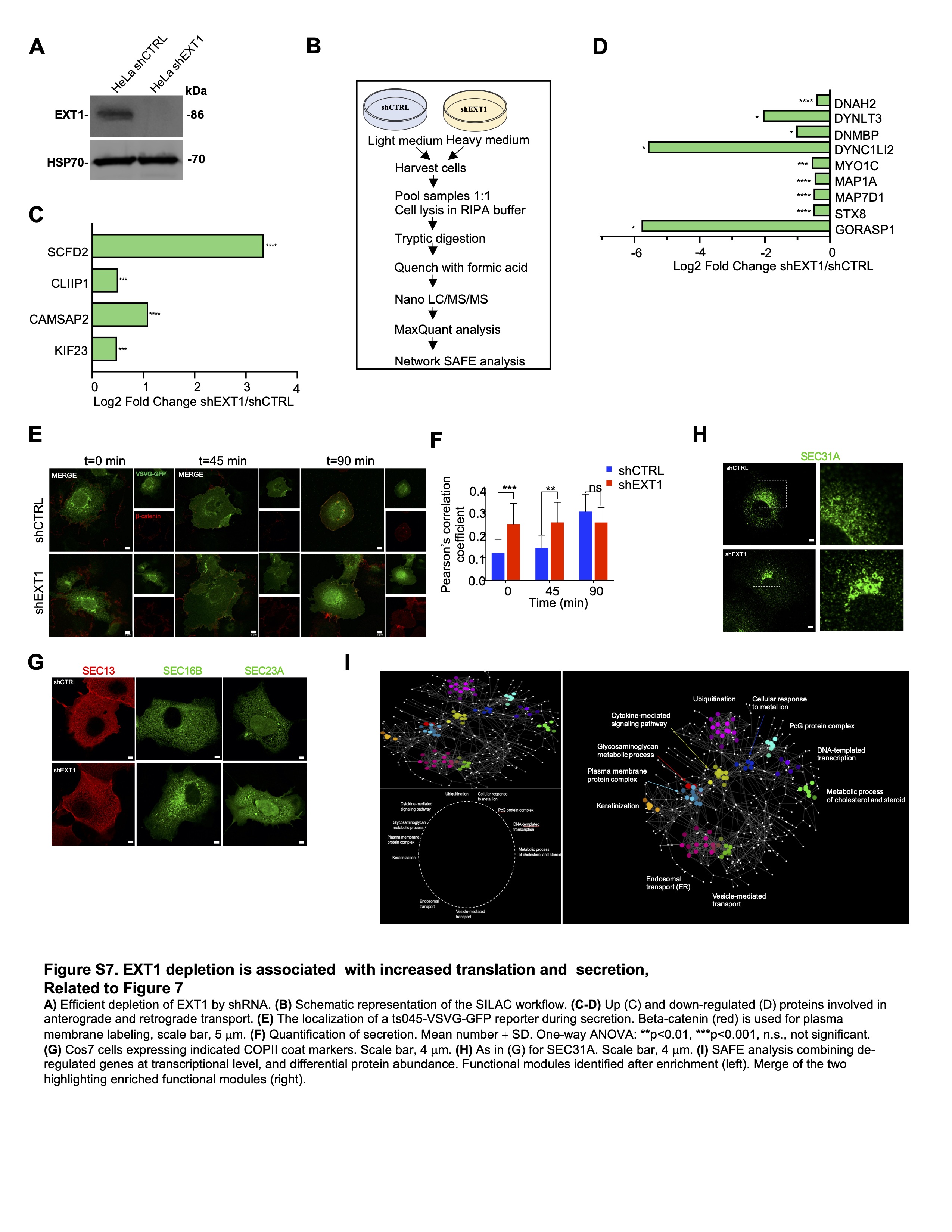
