## Supplemental Figure 3 for "Exostosin-1 Glycosyltransferase Regulates Endoplasmic Reticulum Architecture and Dynamics"

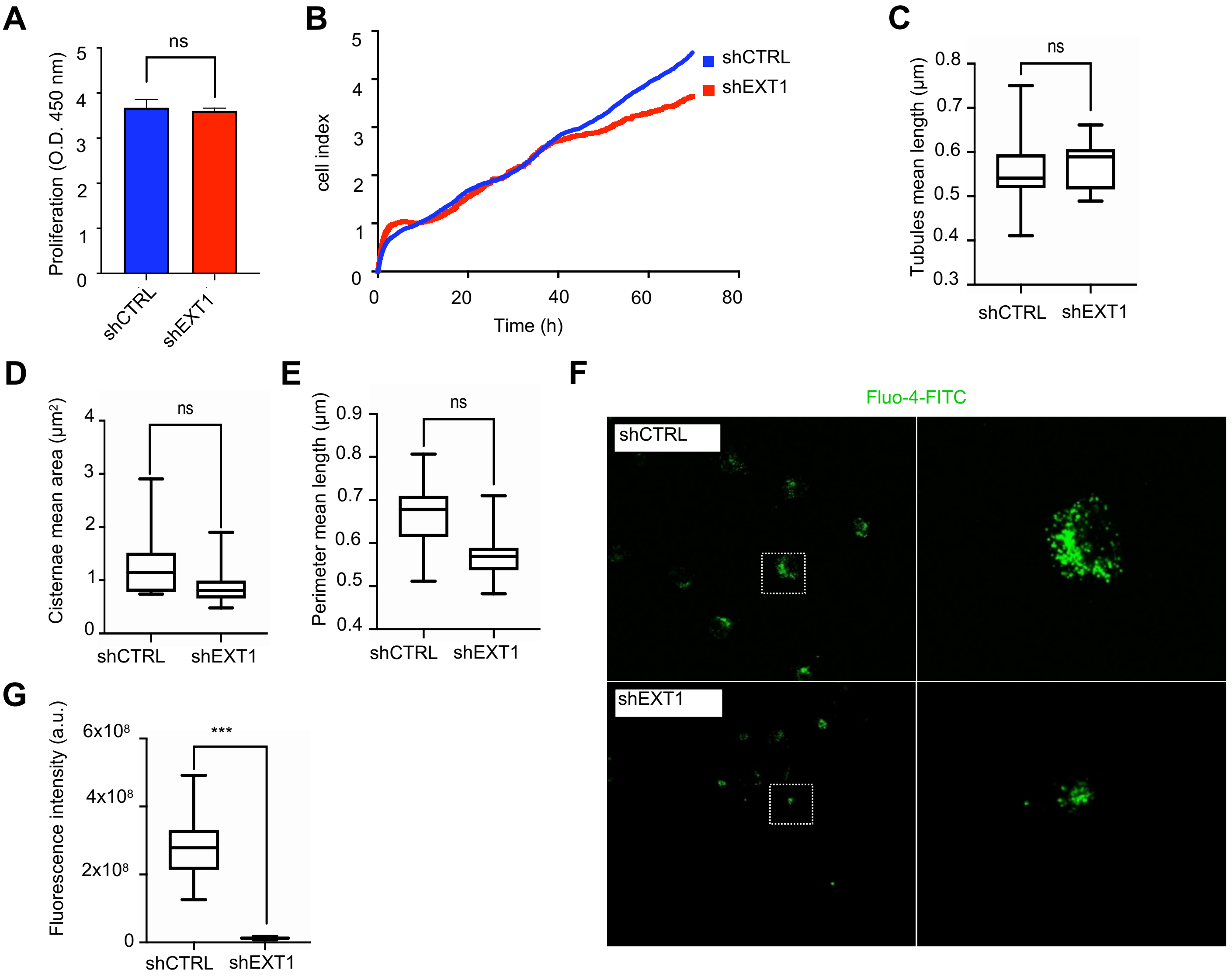


**Figure S3. EXT1 reduction induces ER tubules re-organization and a shift in calcium flux, Related to Figure 3**

**(A)** Proliferation of HeLa cells measured with BrdU optical absorbance at 450 nm. Mean number + SD is plotted. One-way ANOVA: ns: not significant. **(B)** Xcelligence system to compare cell indexes. **(C-E)** Quantitative analysis based on the skeletonization model of Cos7 cells expressing Sec61b. (C) Tubule mean length. Box plot indicates the mean and whiskers show the minimum and maximum values (n = 19-24). (D) As in (C) but for the cisternal mean area. (E) As in (C) but for the perimeter mean length. **(F)** Fluo-4 dye was used to measure Ca2+ concentration in Cos7 shCTRL and shEXT1 cells. **(G)** Mean normalized fluorescence intensity (a.u.) was measured (n = 12). Boxplot indicates the mean and whiskers show the minimum and maximum values. One-way ANOVA: ***p<0.001.
