## Supplemental Figure 4 for "Exostosin-1 Glycosyltransferase Regulates Endoplasmic Reticulum Architecture and Dynamics"

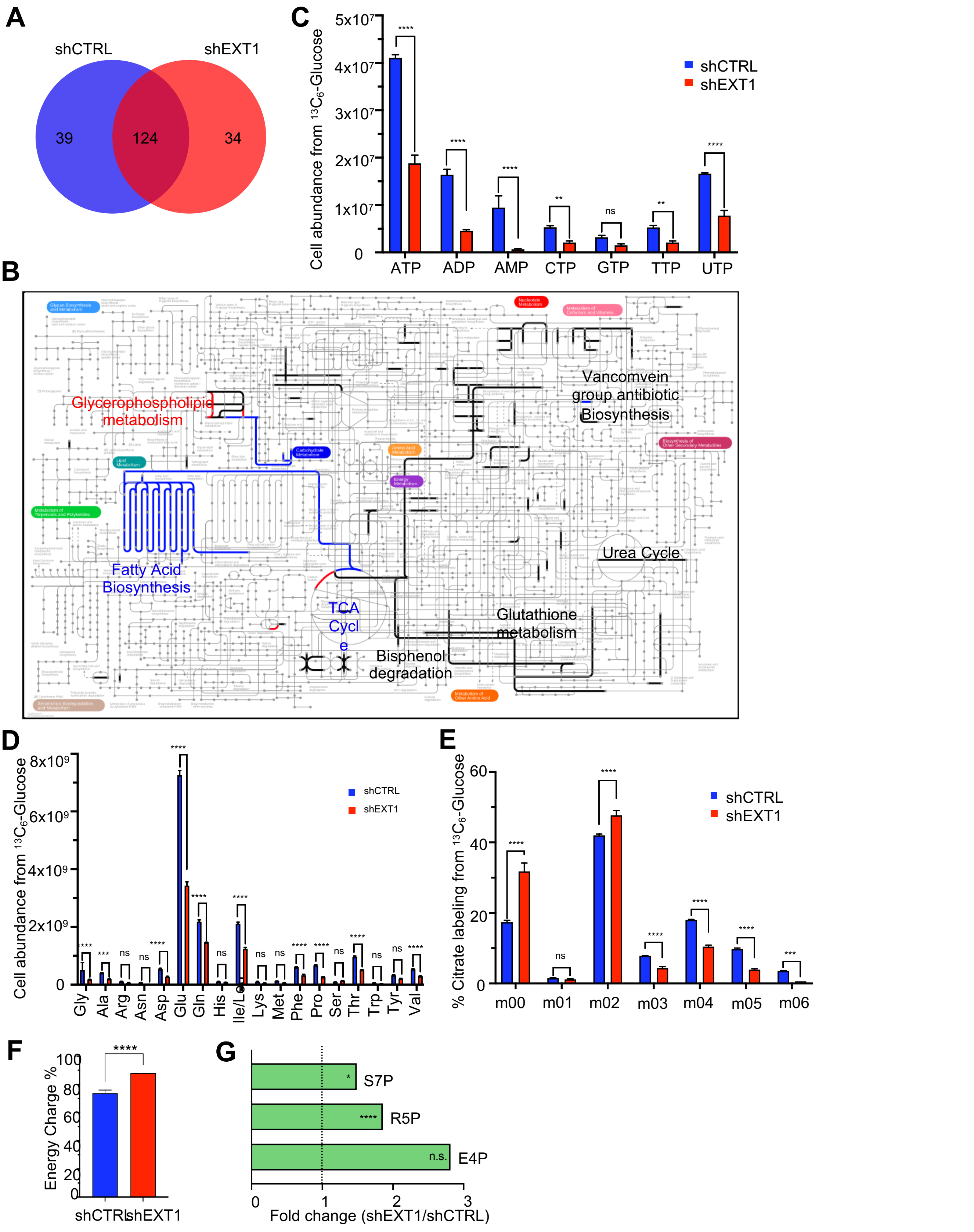


**Figure S4. EXT1 reduction induces a metabolic switch, Related to Figure 4**

**(A)** Venn diagram showing active reactions in control model (blue) and *EXT1* knocked-down model (red). **(B)** Pathways enriched in the active reactions. Blue, Red and black correspond respectively, to reactions uniquely in shCTRL, shEXT1, and in both models. **(C)** Metabolomic analysis from $^{13}C_6$-Glucose of glycolysis nucleotide metabolites in HEK293 cells. Fold change in the abundance of the metabolites in shEXT1/shCTRL is shown. **(D)** As in (C) for amino acid metabolites. **(E)** As in (C) for citrate metabolites. **(F)** Percentage of energy charge. Energy Charge is calculated as (ATP+(0.5*ADP))/SUM(ATP+ADP+AMP), (n = 3). **(G)** As in (C) for pentose phosphate pathway metabolites. One-way ANOVA: *p<0.05, **p<0.01, ***p<0.001, ****p<0.0001, n.s., not significant.
